# Altered chromatin accessibility mediated by somatic cohesin mutations amplifies FLI1-Menin output to promote leukemogenesis

**DOI:** 10.1101/2025.04.01.646632

**Authors:** Jane J. Xu, Viviana Scoca, Yi Chen, Yingqian A. Zhan, Alexander Fisher, Eno-obong Udoh, Sebastian Fernando, Besmira Alija, John Pantazi, Varun Sudunagunta, Edna Stewart, Anabella Maria D. Galang, Michael Williams, Govind Bhagat, Claudia Gebhard, Valeria Visconte, Sarah L Ondrejka, Ruud Delwel, Ming Hu, Richard Koche, Aaron D. Viny

**Affiliations:** Division of Hematology & Oncology, Department of Medicine, Vagelos College of Physicians & Surgeons, Columbia University Irving Medical Center, New York, NY USA; Columbia Stem Cell Initiative, Department of Genetics & Development, Columbia University Irving Medical Center, New York, NY USA; Herbert Irving Comprehensive Cancer Center, Columbia University Irving Medical Center, New York, NY USA; Center for Epigenetic Research, Memorial Sloan Kettering Institute, New York, NY USA; Department of Internal Medicine III, University Hospital Regensburg, Regensburg, Germany; Leibniz Institute for Immunotherapy, Regensburg, Germany; Department of Hematology, Erasmus MC Cancer Institute, Rotterdam, the Netherlands; Department of Translational Hematology and Oncology Research, Cleveland Clinic Lerner College of Medicine, Cleveland, OH USA; Department of Quantitative Health Sciences, Lerner Research Institute, Cleveland Clinic Foundation, Cleveland, OH, USA; Department of Pathology, Vagelos College of Physicians & Surgeons, Columbia University Irving Medical Center, New York, NY USA; Department of Pathology and Laboratory Medicine, Diagnostics Institute, Cleveland Clinic, Cleveland, OH USA; Oncode Institute, Utrecht, the Netherlands

**Keywords:** FLI1, Chromatin architecture, Stem cell, STAG2 leukemia, Revumenib

## Abstract

Chromatin architecture governs transcriptional output and cell identity; however, how its disruption promotes leukemic transformation remains incompletely understood. Here, we show that loss of the cohesin subunit STAG2 creates a hyper-accessible chromatin landscape that amplifies FLI1 activity and extends its binding to ectopic loci. Using multi-omic analyses in human AML samples, cell lines, and mouse models, we identify chromatin-dependent co-occupancy of FLI1 and Menin, accompanied by amplification of Menin occupancy at non-canonical loci beyond its HOXA/MEIS1 targets. Therapeutically, the aberrant expansion of Menin binding drives an altered response towards Revumenib and activates interferon response pathways, creating a therapeutic vulnerability to Menin inhibition. Functionally, STAG2/NPM1c co-mutation drives a stem cell-like immunophenotype and a fully penetrant leukemia *in vivo*. Collectively, these findings define a model in which cohesin loss rewires transcription factor occupancy to amplify oncogenic chromatin programs and expose context-specific therapeutic dependencies.

**Highlights:**

- Chromatin contact density decreases during stem to myeloid differentiation.
- STAG2 loss induces a hyper-accessible chromatin state and expands FLI1 binding.
- FLI1 and Menin co-occupy regulatory loci associated with stemness programs and NPM1c leukemia.
- STAG2/NPM1c co-mutant leukemia exhibit increased Menin occupancy and altered response to Menin inhibition.
- Stag2/Npm1c co-mutation drives a dysplastic, transplantable AML.

## Introduction

Hematopoiesis requires a tightly regulated balance between self-renewal and differentiation of stem cells to produce lineage-committed progeny, a process disrupted during leukemogenesis ^1,2^. Hematopoietic stem cells (HSCs) are multipotent cells that can give rise to all blood cell types required to sustain hematopoiesis^3,4^. The molecular mechanisms that regulate HSC fate decisions and lineage specification are of great interest for understanding hematopoiesis and its dysregulation in diseases. Chromatin dynamics during hematopoiesis reflect the complex interplay between epigenetic modifications ^5,6^, chromatin remodeling ^7,8^, and transcription factors (TFs) binding at cis-regulatory elements (CREs) ^9,10^ that control gene expression^11^. Among these TFs, several, including the ETS-family member FLI1 play a crucial role in HSC function and self-renewal, regulating key hematopoietic genes through specific chromatin interactions.

Cohesin and CTCF are key regulators of chromatin architecture, forming loops at the boundaries of topologically associated domains (TADs) to insulate transcriptional programs^12^. Despite their known roles in genome organization, the temporal and spatial chromatin changes that occur during HSC differentiation remain incompletely understood. In acute myeloid leukemia (AML), aberrant transcriptional activity, DNA methylation, and 3D chromatin disorganization have been well documented^13–15^. Aberrant chromatin regulation in AML is also propagated through menin, which scaffolds the KMT2A (MLL) complex at HOXA loci to sustain the transcriptional program required for leukemic stem cell maintenance^16^. Notably, cohesin mutations occur in ∼12% of AML and ∼20% of high-risk myelodysplastic syndromes (MDS)^17,18^, with mutations in the subunit STAG2 representing the majority of cases^19^.

STAG2 mutations are predicted to arise early in clonal evolution and are often co-mutated with other AML-defining lesions such as NPM1c, RUNX1, and SRSF2^17,20,21^. Clinically, STAG2 is a defining mutation for AML with myelodysplasia-related changes (AML-MRC), a subtype with distinct molecular features and prognosis^22,23^. We and others have shown that STAG2 loss leads to expansion of hematopoietic stem and progenitor cells (HSPCs) and impaired lineage commitment^24–27^. While cohesin mutations disrupt chromatin architecture and accessibility— potentially affecting transcription factor binding patterns and target gene expression in leukemic cells—the specific mechanisms by which STAG2 mutations drive leukemic transformation remain unclear. Although previous work has generated co-mutational Smc3 leukemia murine models of AML, no experimental model exists for STAG2-mutant AML, the most frequent cohesin mutation in AML^19,28,29^. Importantly, co-mutation patterns differ substantially between de novo and secondary AML^21^, particularly for mutations associated with antecedent myelodysplasia^30^. We chose to investigate the NPM1c co-mutant setting not because of preferential co-mutation with STAG2, but because NPM1c leukemia retains a well-defined Menin dependency that provides a tractable model to interrogate chromatin-dependent transcriptional regulation.

In this study, we investigated chromatin alterations during the LSK to GMP transition and examined the effects of co-STAG2/NPM1c mutation on chromatin accessibility and Menin engagement across human cells, murine models, and AML patient samples. We identify a conserved hyper-accessible chromatin state following STAG2 loss, associated with ectopic FLI1 binding and activation of stemness-associated programs, including CD34 expression. Consistent with these findings, STAG2/NPM1c co-mutant AML exhibits a CD34+ immunophenotype in patients. Functionally, co-mutation of Stag2 and Npm1c drives a fully penetrant AML with myelodysplastic features arising from myeloid-biased HSPCs. Mechanistically, STAG2 loss promotes ectopic FLI1 binding, that is associated with the expansion of Menin occupancy at non-canonical loci, enhancing sensitivity to Menin inhibition. Collectively, these findings define a chromatin-driven mechanism of leukemogenesis whereby aberrant architecture can co-opt endogenous TF signaling and reveal a targetable dependency in STAG2-mutant AML.

## Results

### Chromatin contact dissolution characterizes the HSPC-to-myeloid transition

To define dynamic chromatin remodeling events during hematopoietic differentiation, we compared stem cell–enriched LSK cells (Lineage-Sca1+cKit+) to committed granulocyte– macrophage progenitors (GMP; Lineage-cKit+Sca1-Cd34+FcR *γ* +). We applied a multi-omic approach to compare these functionally distinct populations within the hematopoietic hierarchy. Using 3D genome organization and chromatin accessibility techniques, we defined the epigenomic changes during the transition from LSK to GMP in wild-type (WT) mice and then determined the impact of Stag2 loss on this process. Previously, we demonstrated *Stag2* loss leads to expansion and alters lineage commitment of HSPCs **(Supplemental Figure 1A-C)** ^27^, however the 3D genome re-organization and the role of Stag2 during myeloid commitment have not yet been identified.

To characterize the 3D chromatin structure during LSK to GMP transition, we performed low-input Hi-C on LSK and GMP population (n=3 biological replicates for each cell type) (**Figure 1A**). Globally, we observed changes at the compartment, TADs, and loop level when comparing the LSK and GMP cells. We found that there are more long-range contacts (>10Mb) (**Figure 1B**), higher TAD (50Kb bin) insulation **(Supplemental Figure 1D)**. These data indicate that hematopoietic stem cells exhibit a highly interconnected chromatin architecture that is progressively pruned during transition to lineage-committed progenitors.

**Figure 1:**
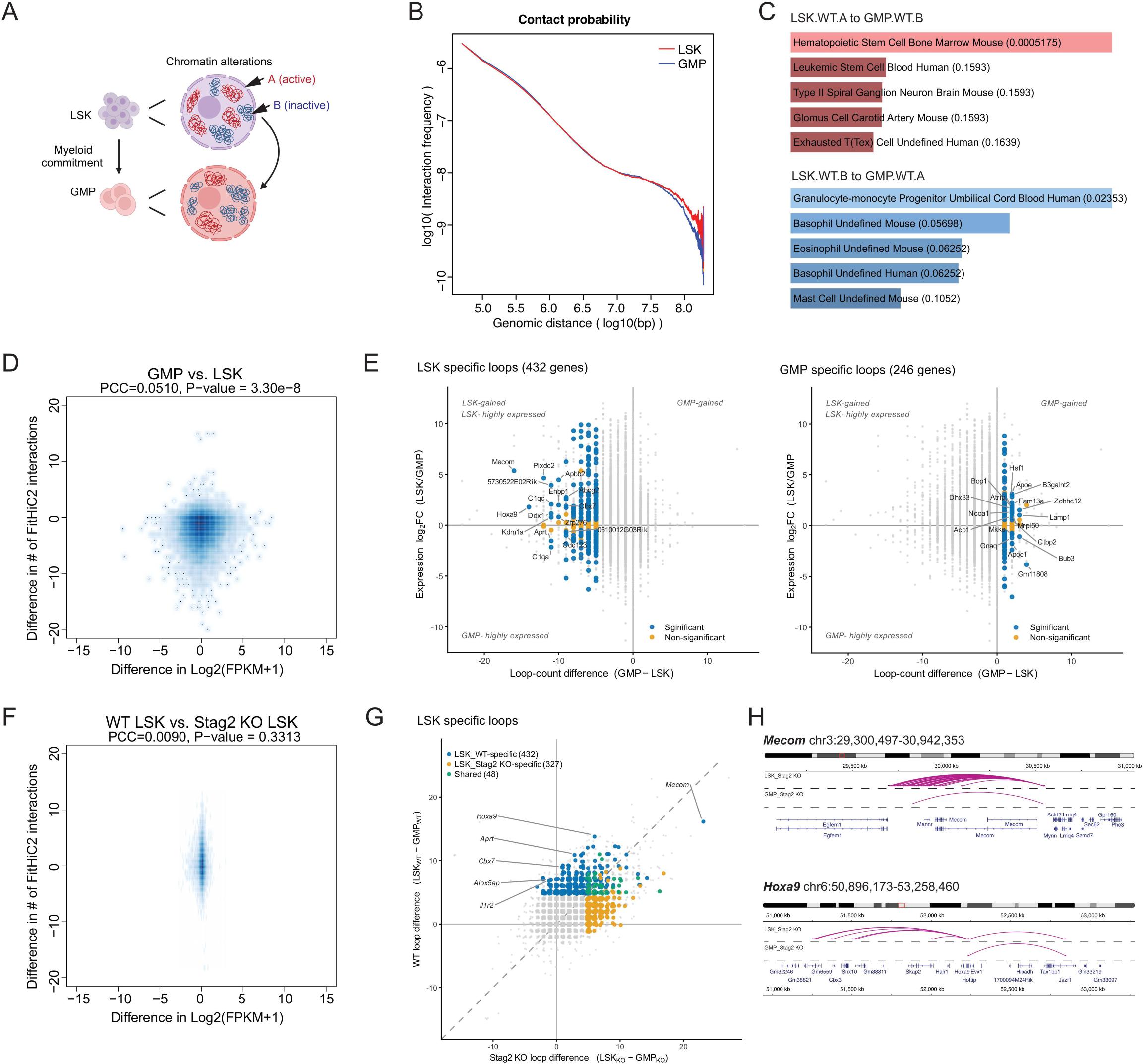
Pruning of chromatin contact during the hematopoietic stem and progenitors to myeloid transition. **A)** Schematic of LSK to GMP transition and chromatin compartments. **B)** The contact probability plot of LSK and GMP cells. Pooled data from 3 biological replicates. **C)** The cell type annotation atlas analysis of A/B compartment genes in LSK and GMP cells. **D)** Heatmap of differentially expressed genes intersect with differential enhancer promoter interactions (FitHiC2) in GMP vs LSK. Cut-off for the differentially expressed genes are log Fold Change (logFC) > log2(1.5), False Discovery Rate (FDR) < 0.1. **E)** Per-gene HiC loop-count difference versus gene expression changes for genes with detected loops in LSK and GMP. The LSK specific loops (432 genes) are highlighted in the left panel, and GMP specific loops (246 gene s) are highlighted in the right panel, colored by differential expression (blue, FDR < 0.05; orange, n.s.; Benjamini–Hochberg), and other genes are colored grey. Selected LSK loop-gained genes are labeled. n = 10,572 genes. **F)** Heatmap of differentially expressed genes intersect with differential enhancer promoter interactions (FitHiC2) in WT LSK vs Stag2Δ LSK. Cut-off for the differentially expressed genes are log Fold Change (logFC) > log2(1.5), False Discovery Rate (FDR) < 0.1. **G)** Scatter plot comparing differential chromatin looping per-gene across the LSK to GMP transition in each genotype. The LSK WT specific loops are labelled in blue, Stag2Δ LSK specific loops are labelled in orange, the shared loops in blue. The non-LSK specific loops are in grey. H) Lineplot of Mecom loops (upper panel) and Hoxa9 loops (lower panel) in Stag2Δ LSK and Stag2Δ GMP cells.

We next set out to determine if the 3D genome reorganization is associated with cell type shift and A/B compartment changes during stem to myeloid differentiation. Compartment A represents accessible chromatin with higher gene expression, while chromatin in compartment B is largely compacted with repressive histone mark and few gene activities^12,31,32^, In our analysis, we identified 137 genes that were in the inactive (B) compartment of LSK cells and then shifted to the active (A) compartment in GMP cells, which we labelled as LSK.B to GMP.A. On the contrast, there were 109 genes that moved from active compartment of LSK cells to inactive compartment of GMP cells, which we labelled as LSK.A to GMP.B. Using the cell annotation atlas^33^, genes that changed from LSK.B to GMP.A are correlated with ‘Granulocyte-monocyte Progenitor’ cell type, where genes changed from LSK.A to GMP.B are associated with a cell type of ‘hematopoietic stem cell’ (**Figure 1C**). This suggests that the A /B compartment switch during the LSK to GMP transition is correlated with cell fate, as described in other tissue type^34^.

At loop level, we observed more enhancer-promoter (E-P) interactions (10Kb FitHiC2 interactions) in the LSK cells, compared to GMP cells (**Figure 1D, Supplemental Table 1)**. In total, we identified 12,680 loops that remain stable during the transition, whereas 43,482 loops are LSK-specific, and 10,553 loops are GMP-specific (FitHiC2 loops, 10Kb bins). This corresponds to 432 genes with high frequency contacts (FDR<1%) in LSK and no loop in GMP. Consistent with this, stem-associated loci such as Mecom and Hoxa9 exhibited reduced looping and decreased expression during differentiation, illustrating coordinated loss of regulatory interactions at key stemness genes (**Figure 1E, Supplemental Figure 1E-F)**. On the other hand, we observed only 246 genes with GMP-specific loops, and key myeloid genes such as *Ms4a3* were absent from this set (**Figure 1F, Supplemental Figure 1G, Supplemental Table 2)**. These data suggest that dynamic chromatin interactions are more closely associated with regulation of stemness rather than the downstream myeloid progenitor.

We next examined whether STAG2 loss disrupts the chromatin reorganization that accompanies myeloid differentiation. Consistent with our prior work^27^ and that of others^20,35,36^, no global changes in TAD insulation were observed with Stag2 loss in LSK or GMP cells when within the same cell type **(Supplemental Figure 1H)**. Similarly, we found similar trend of chromatin dissolution when LSK exits stemness (**Supplemental Table 1).** However, when Stag2 is knocked out, the number of genes with high frequency contacts (≥5 loops) specific to LSK drops from 432 to 327, while genes with GMP-specific high frequency contact loops increases from 246 to 600 **(Supplemental Table 2)**. At loop level, we observed that deletion of Stag2 in the LSK population did not significantly alter E-P interactions between WT LSK and Stag2 mutant (KO) LSK cells (**Figure 1G**). However, Stag2 loss impaired the specificity of chromatin interactions, leading to persistence and mislocalization of loops at stem-associated loci. For example, stem-associated loci such as *Mecom* and *Hoxa9* exhibited persistent chromatin interactions in Stag2-deficient cells, despite differentiation (**Figure 1H**). To determine if disruption of the looping activities observed in the Stag2 mutant LSK cells will affect the genes expression program, we selected two pathways that are significantly enriched for LSK or GMP cells respectively, which are hematopoiesis stem cell long term or myeloid Cebpα network, based on the Gene Set Enrichment Analysis (GSEA) analysis. We found that Stag2 mutant LSK cells have more stemness and increased myeloid activity comparing to LSK WT **(Supplemental Figure 1I, Supplemental Table 3)**. Consistent with altered chromatin architecture, Stag2-deficient LSK cells display activation of stemness-associated transcriptional programs alongside aberrant myeloid gene activity.

Collectively, these findings demonstrate that hematopoietic differentiation is accompanied by extensive pruning of chromatin contacts and regulatory interactions. Loss of STAG2 disrupts this organization, resulting in persistence and mislocalization of stem-associated chromatin interactions and gene expression programs. These findings raise the possibility that aberrant chromatin organization in STAG2-deficient cells may alter transcription factor access to regulatory elements, thereby reshaping downstream transcriptional programs.

### STAG2 loss amplifies chromatin accessibility in stem and leukemic cells

After evaluating the 3D genome organization changes, we performed bulk ATACseq on WT LSK and GMP populations (n=3 biological replicates for each cell type) to assess the chromatin accessibility change during stem to myeloid fate commitment (**Figure 2A**). We identified 50,404 dynamically accessible regions which were nearly balanced for gains and losses in accessibility **(Supplemental Figure 1J, upper panel)**. Using HOMER motif enrichment analysis, we identified that WT LSK cells exhibit open and accessible chromatin at Fli1-motif, with closure of these regions upon differentiation towards a GMP cell state. By contrast, the WT GMP cells increased accessibility at Ctcf targets (**Figure 2B, Supplemental Table 4&5A)**. We found that *Stag2* KO LSK cells have increased accessibility, especially at motif of Ews:Fli1 and Fli1, without changes at downstream pathway (**Figure 2C, Supplemental Figure 1J lower panel, Supplemental Table 5B)**. While extensive chromatin changes were observed between Stag2 KO and WT cells, we found no differential expression changes of any of the hematopoietic “heptad TFs” (Fli1, Erg, Gata2, Runx1, Tal1, Lyl1, Lmo2), which are key regulators of normal and leukemic hematopoiesis^37^ **(Supplemental Figure 1K)**. To validate our findings in human cells, we performed siRNA knockdown of STAG2 *in vitro* in human CD34+ cells, which contains the most primitive hematopoietic stem and progenitor cells and proceeded with ATACseq (**Figure 2D**). We observed both increased and decreased chromatin accessibility (**Figure 2E**). Again, we identified FLI1 as the top motif gaining accessibility, which confirms our observation in murine LSK cells. This suggests that STAG2 loss alters accessibility of the chromatin which could affect TF binding landscapes without changing TF abundance, highlighting a chromatin accessibility-driven mechanism that potentiates TF activity in STAG2 mutants.

**Figure 2:**
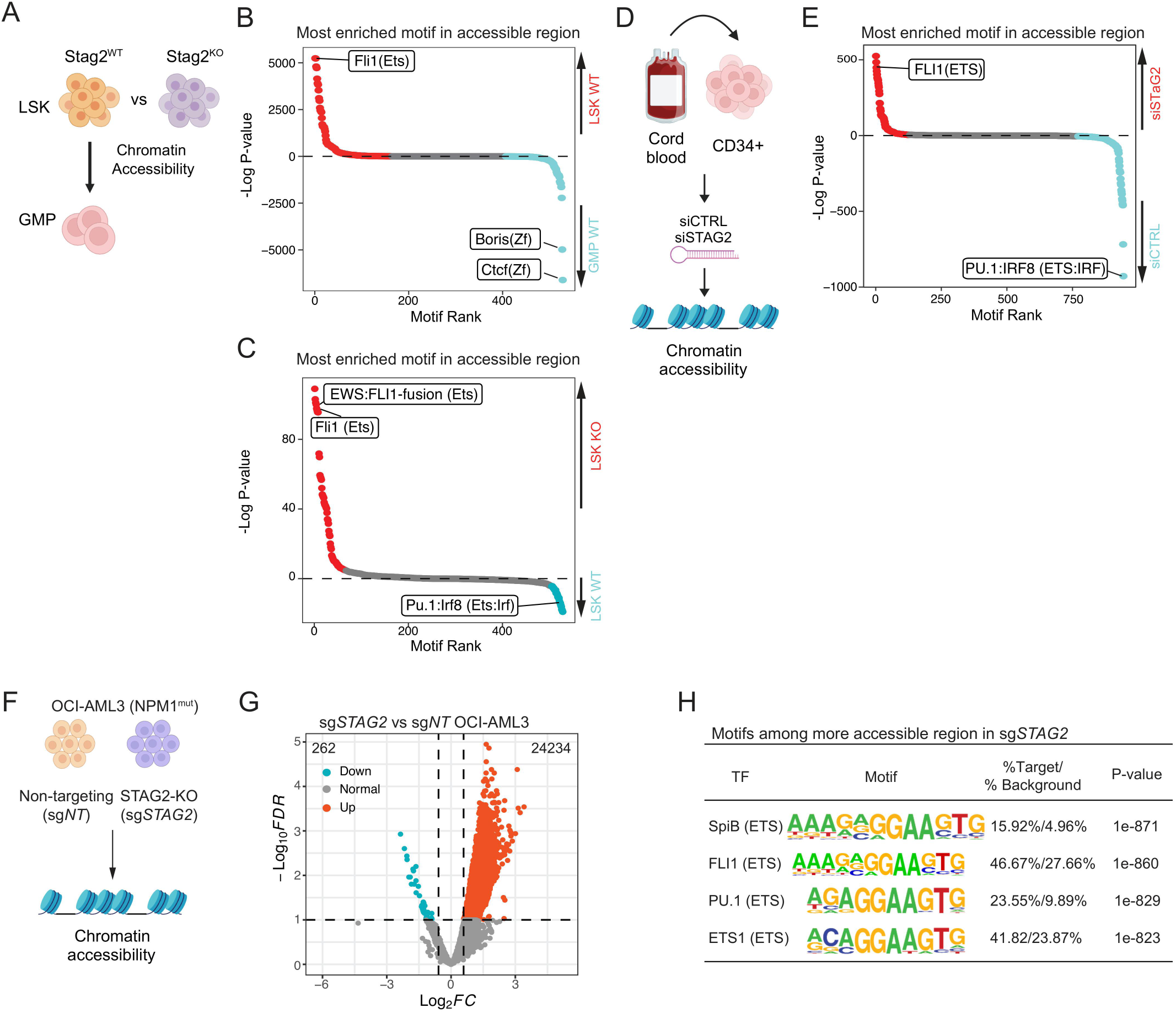
STAG2 loss increases chromatin accessibility in HSPC and leukemia. **A)** Schematic of chromatin accessibility comparison of LSK vs GMP, and WT LSK vs Stag2Δ (KO) LSK. **B)** Motif enrichment comparison of chromatin accessibility between WT LSK and WT GMP cells (n=3 biological replicates). **C)** The motif enrichment comparison between WT LSK and Stag2Δ (KO) LSK cells (n=3 biological replicates). **D)** Schematic of siRNA knockdown and ATAC-seq on CD34+ cells. **E)** Motif enrichment comparison of chromatin accessibility between siCTRL and siSTAG2. **F)** Schematic of chromatin accessibility comparison between isogenic sg*NT* and sg*STAG2* OCI-AML3 cells. **G)** The chromatin accessibility changes in the sg*STAG2* compared to the sg*NT* cells (n=2 independent clones per genotype). **H)** The Known Homer motif analysis of more accessible sites in the sg*STAG2* cells. For listed TFs, p-value as labelled and q value are 0 (BH-adjusted).

Next, we sought to determine if STAG2 loss also leads to chromatin accessibility change in leukemia. NPM1c is one of the most common AML mutations and results in chromatin binding of NPM1c at the *HOXA* locus and *MEIS1* resulting in leukemogenesis ^38,39^. STAG2 mutation co-occur with NPM1c and is consistent with a myelodysplasia-related disease context prior to AML formation, which is associated with altered therapy response and prognosis ^21^. Therefore, we chose the OCI-AML3, which is a NPM1c mutant AML cell line, and generated isogenic single cell-derived CRISPR non-targeting (sg*NT*) and knock-out (sg*STAG2*) clones to model cohesin mutant leukemia (**Figure 2F, Supplemental Figure 2A-B)**. This modeling strategy reflects clinically observed mutational states in which myelodysplasia-associated lesions coexist with AML-defining drivers such as NPM1c and enabled us to directly assess the impact of STAG2 loss on chromatin accessibility within an NPM1c-driven leukemic context.

In liquid culture, sg*STAG2* clones displayed a decreased expansion rate compared to sg*NT* **(Supplemental Figure 2C)** which was corroborated by a lower percentage of cells in S phase as measured by BrdU incorporation **(Supplemental Figure 2D)**. Through ATAC-seq, we observed a marked increase in overall chromatin accessibility in sg*STAG2* cells with 24,496 differentially accessible peaks (**Figure 2G**). These sg*STAG2*-specific peaks were enriched for targets of multiple ETS family TFs, including FLI1, which represented 46.6% of targets (**Figure 2H**). These data suggest a STAG2-driven chromatin regulatory mechanism centered on the transcription factor FLI1 in both mouse and human hematopoietic stem and progenitor cells and conserved in STAG2 mutant human leukemia.

### STAG2 loss drives ectopic FLI1 binding and amplifies the Menin cistrome in NPM1c leukemia

FLI1, an ETS family transcription factor, is an essential hematopoietic TFs and important in stem cell self-renewal, regulating many key factors including *MECOM* ^37^. To determine the potential for the aberrant accessibility of FLI1 targets in contributing towards hematopoietic transformation in STAG2 mutation, we interrogated published AML datasets and found that FLI1 targets were strongly enriched for NPM1c binding ^40,41^. Therefore, we hypothesized that STAG2 loss increases accessibility at FLI1 binding sites and cooperates with the NPM1c related chromatin factors to facilitate transformation.

As ETS family TFs have similar motifs, we assessed the FLI1-specific contribution through cell fractionated western blotting and chromatin immunocapture using CUT&RUN. In addition, we profiled chromatin binding of Menin through CUT&RUN, as Menin is an oncogenic partner and therapeutic target in NPM1c AML^42^. Although no difference in overall or cytoplasmic protein expression was observed, the *sgSTAG2* cells have a specific increase in the chromatin-bound FLI1 and Menin through western blotting **(Supplemental Figure 2E)**. Consistent with this data, we observed a marked increase in FLI1, as well as Menin peaks by CUT&RUN in *sgSTAG2* OCI-AML3 cells (**Figure 3A-B, Supplemental Table 6 & 7)**. We found that most FLI1 targets coincide with sites of increased chromatin accessibility in sg*STAG2* cells and show higher H3K27ac signal, which marks active promoters and enhancers **(Supplemental Figure 2F, Supplemental Table 8)**. To test whether ectopic FLI1 binding mirrors how FLI1 normally engages chromatin, we compared the FLI1 peaks in sg*STAG2* cells with ChIP-seq data generated human CD34+ cells ^43^ and found over one-third of FLI1 binding sites in STAG2-deficient AML cells overlap with those in normal human HSPCs, including at stem cell marker genes, *CD34* **(Supplemental Figure 2G**, **Supplemental Table 8**). This data suggests these FLI1 binding partially resembles the existing binding pattern in the normal HSPCs. To determine the impact of FLI1 binding on gene expression, we performed transcriptomic analysis and found 641 genes with decreased expression in sg*STAG2*, and 392 genes with increased expression compared to sg*NT* **(Supplemental Figure 2H)**. Further characterization found FLI1 binding sites on the promoter region of 165 upregulated and 306 down-regulated genes, respectively **(Supplemental Figure 2I, Supplemental Table 9a&b)**.

**Figure 3:**
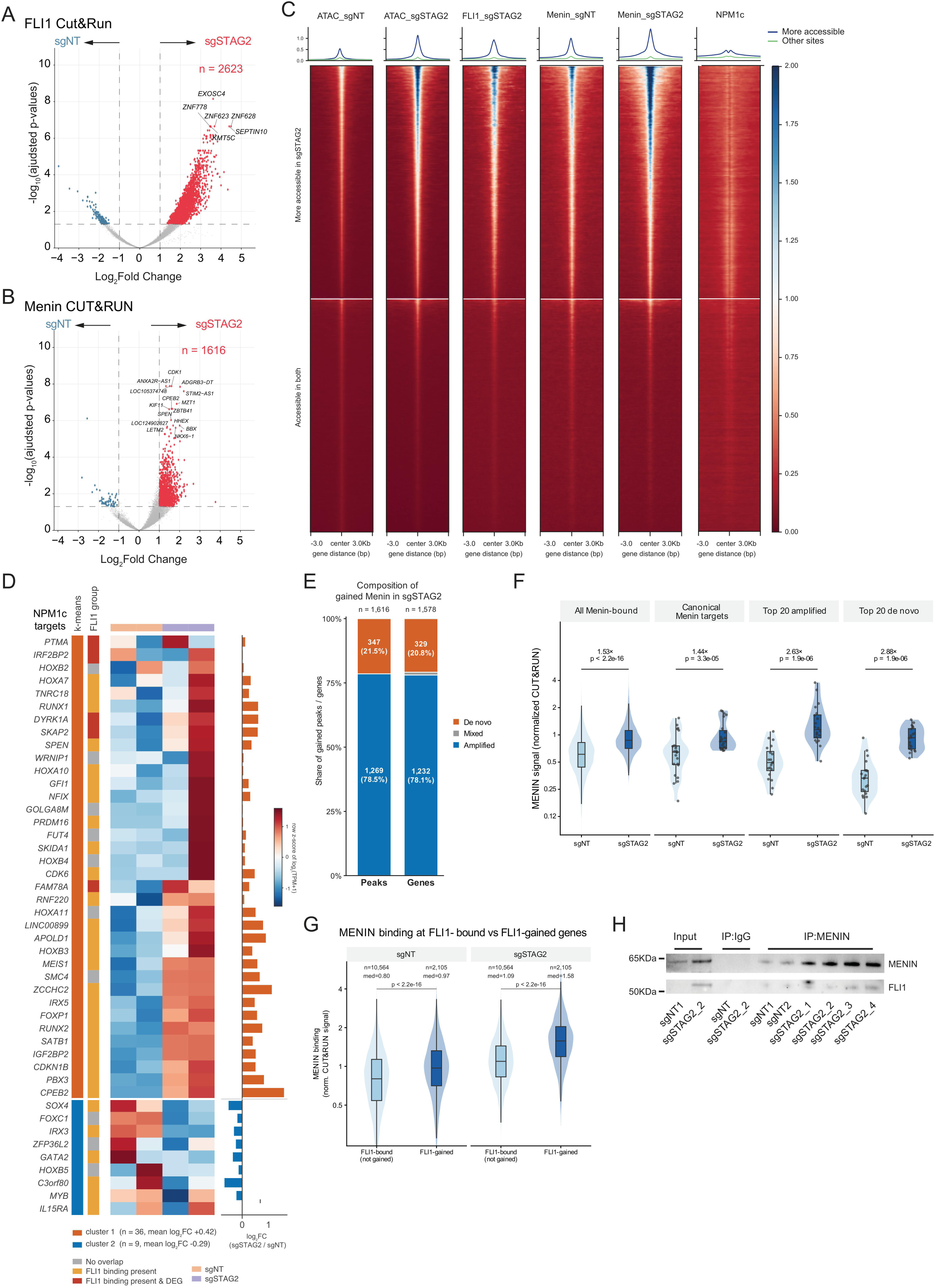
STAG2 loss drives ectopic FLI1 binding and amplifies Menin cistromes in leukemia. **A)** Volcano plot of differential FLI1 binding sites in sg*STAG2* vs sg*NT*. Significant binding is labelled in red sg*STAG2* and blue for sg*NT*, padj < 0.05 and log2FC > 1. **B)** Volcano plot of differential Menin binding sites in sg*STAG2* vs sg*NT*. Significant binding is labelled in red sg*STAG2* and blue for sg*NT*, padj < 0.05 and log2FC > 1. **C)** Heatmap of chromatin accessibility, FLI1, Menin and NPM1c Cut&Run track grouped by more accessible region in sg*STAG2* and other sites. **D)** K-mean based Heatmap (k=2) showing the number of NPM1c target genes that have FLI1 binding in the sg*STAG2*cells (yellow) and have differential gene expression (red). No binding genes are in grey. Log2 fold-change of gene expression is on the right of the heatmap. **E)** Barplot of de novo or amplified Menin binding sites among all sites that gained Menin signal in the sg*STAG2* cells. Gained Menin signal is defined as padj < 0.05 and log2FoldChange > 1. **F)** Menin signal at canonical and gained target sets, including all Menin-bound genes, canonical Menin targets, the top 20 amplified gained and the top 20 de novo gained genes, in sg*NT* versus sg*STAG2*. Median fold-change of sg*STAG2*/sg*NT* labelled on top. Paired two-sided Wilcoxon test. **G)** Comparison of Menin signal at FLI1 bound or FLI1 gained genes in sg*NT* and sg*STAG2*. Wilcoxon rank-sum test. **H)** Co-Immunoprecipitation of Menin in sg*NT* and sg*STAG2* chromatin fraction, blotted for Menin and FLI1.

Based on our results, we further hypothesize that FLI1 could partner with existing chromatin binding factors in NPM1c leukemia, such as NPM1c and Menin, to drive leukemic program. To investigate the relationship between ectopic FLI1 binding, NPM1c, and Menin occupancy, we compare the binding profile of FLI1, Menin from sgSTAG2 OCI-AML3 cells with published NPM1c peaks^40^. Comparing all three factors, we found that FLI1 is co-localized with Menin and NPM1c peaks at the more accessibility region of sg*STAG2* cells. (**Figure 3C, Supplemental Table 10)**. Between FLI1 and NPM1c, FLI1 is bound to 34 of the 44 NPM1c target genes^41^ in the sgSTAG2 and increased the expression of a subset of genes, such as *FOXP1*, *RUNX1* and *PBX3* (**Figure 3D**).

To define the relationship between FLI1 occupancy and Menin localization, we began by quantifying genome-wide Menin occupancy in sg*STAG2* cells. Menin binding was elevated in sg*STAG2* cells relative to sg*NT* controls (**Figure 3C, Supplemental Table 7)**. To distinguish amplification of pre-existing sites from de novo binding, we performed differential binding analysis with DESeq2. Amplified sites were defined as sg*NT* peaks with significantly increased signal in sg*STAG2* (padj < 0.05 & log2FC > 1), and de novo sites as gained peaks (padj < 0.05 & log2FC > 1) not overlapping any sg*NT* -called Menin peak **(Supplemental Table 11)**. In total, increased occupancy was predominantly driven by amplification of existing sites (1,269 peaks / 1,232 genes), with de novo binding representing a smaller fraction (347 peaks / 329 genes) at lower occupancy (median sg*STAG2* signal 1.09 vs. 1.45 log2(CPM+1) at amplified sites) (**Figure 3E**). The amplification magnitude is similar at canonical Menin targets and other Menin-bound genes in the sg*STAG2* (**Figure 3F**). Together, these data suggest that STAG2 loss amplifies the Menin cistromes in OCI-AML3 cells.

Following this, we compared binding profile of FLI1 and Menin in sg*STAG2*cells. Within regions co-bound by FLI1 and Menin, their peak signals were positively correlated between the two factors **(Supplemental Figure 2J)**. Importantly, Menin signal is enriched at FLI1 binding sites in the sg*STAG2* (**Figure 3G**). This is supported by an observation that FLI1 and Menin exhibit chromatin-dependent co-occupancy in sg*STAG2* cells (**Figure 3H).** Notably, reciprocal immunoprecipitation using FLI1 did not recover Menin, indicating that this is not a stable protein– protein interaction but instead reflects chromatin-dependent co-occupancy **(Supplemental Figure 2K)**. Overall, these data demonstrate substantial co-localization of ectopic FLI1 binding with NPM1c-associated and Menin-associated chromatin regions.

### FLI1 binding drives up-regulation of stem cell programs in STAG2-mutant/NPM1c leukemia

Recent studies have demonstrated that AML with cohesin mutations, including STAG2 loss of function mutations, were shown to be highly indicative of a history of myelodysplasia and most frequently presented in the disease category of AML-MRC^21^. We sought to examine if there is a transcriptomic difference between STAG2 mutant AML and non-STAG2 mutant AML. Therefore, we performed analysis on RNAseq data of AML patients with STAG2 mutations without NPM1c, NPM1c mutations without STAG2, co-mutant STAG2 and NPM1c patients, and patients without either mutation (Control). GSEA showed that the co-STAG2/NPM1c mutant AML has an increased gene signature of hematopoietic stem cell program, decreased myeloid cell development and less AML with NPM1c signature compared to single NPM1c mutant AML (**Figure 4B, Supplemental Table 12)**. While we observed no difference in *HOXA9* and *MEIS1* expression, *HLF* levels were decreased in the double mutant AML compared to NPM1c AML **(Supplemental Figure 3A)**. While the pathway analysis suggests that there is a trend of more stemness feature towards co-STAG2/NPM1c AML, we next examined whether the double mutant AML also has a more immunophenotypically stem population. Firstly, in the co-STAG2/NPM1c mutant samples, we found that there is an upward trend of CD34 expression in the double mutant, compared to NPM1c AML (**Figure 4A**).Secondly, to validate CD34 level at the protein level, we examined cell surface markers from clinical flow data from patients with co-mutant STAG2 and NPM1c without STAG2. Similar to prior reports ^44,45^, the NPM1c AML patients overwhelmingly had a CD34 negative immunophenotype, whereas all but one co-mutant STAG2/NPM1c AML patient was CD34-positive (**Figure 4C**) with increased CD34 mean fluorescent intensity by flow cytometry in the blast cells (**Figure 4D, Supplemental Figure 3B)**.

**Figure 4:**
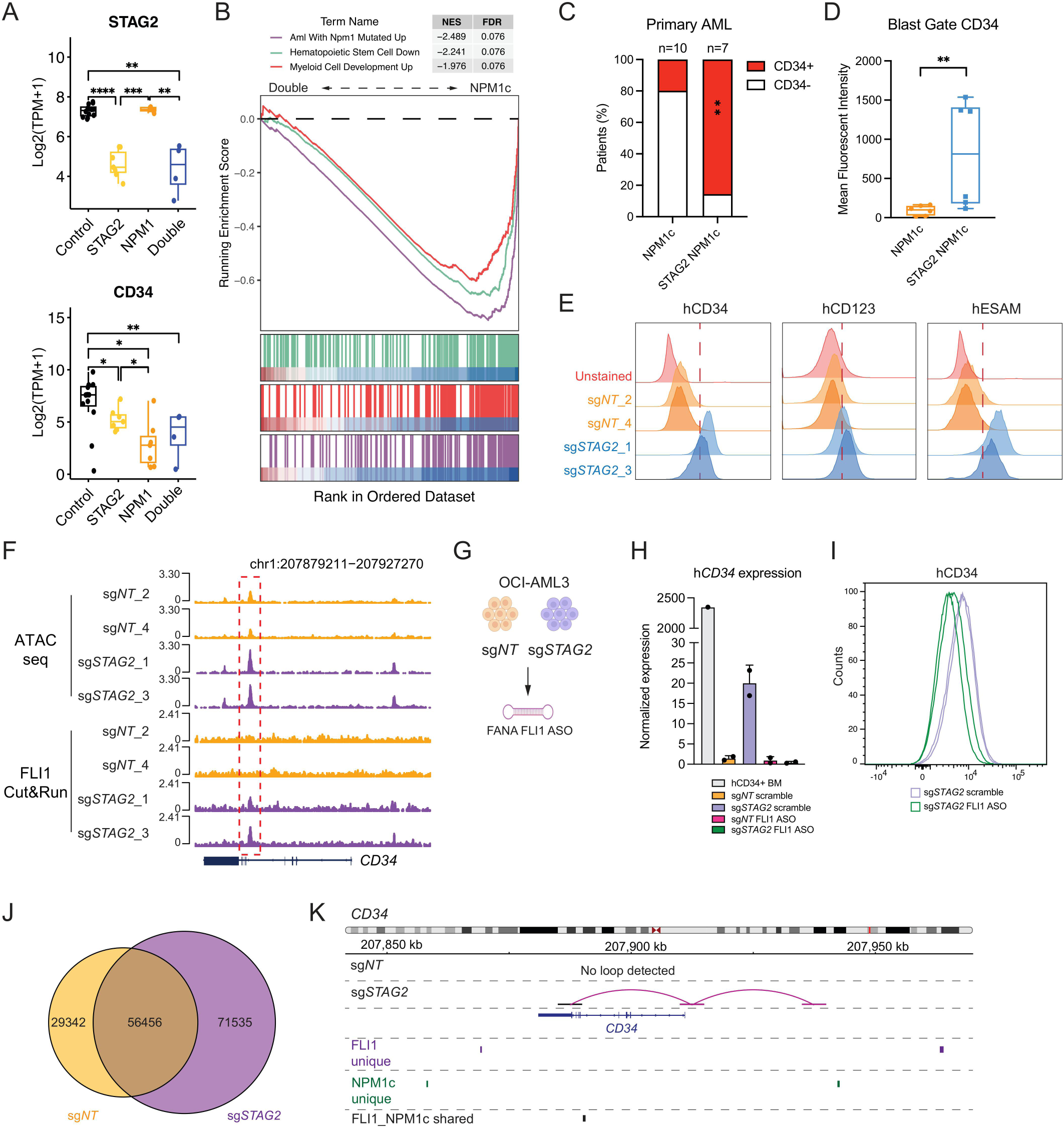
FLI1 binding drives up-regulation of the human stem cell marker CD34 in leukemia. **A)** Boxplot showing STAG2 expression (Top panel) and CD34 expression (Bottom panel) level in control, STAG2 mutant, NPM1c mutant and STAG2 NPM1c mutant AML patients. *p ≤0.05, **p ≤0.01, ***p ≤0.001, ****p ≤0.0001, Wilcoxon signed rank test. **B)** GSEA analysis of STAG2 NPM1c (Double) AML shows increased enrichment of stem cell pathway, and decreased myeloid cell development comparing to NPM1c AML transcriptome. **C-D)** Immunophenotyping of CD34 level in NPM1c and STAG2 NPM1c patient sample showing proportion of CD34 positive patients (C, **p ≤0.01, Chi-square test) and Mean Fluorescent Intensity (MFI) under blast gates (D, **p ≤0.01, unpaired t-test). **E)** Immunophenotyping of CD34, CD123 and ESAM in sg*NT* and sg*STAG2* OCI-AML3 cells. Representative FACS plots of n=3 independent experiments. **F)** IGV track view of the CD34 locus with ATACseq and FLI1 CUT&RUN of sg*NT* and sg*STAG2* OCI-AML3 cells. **G)** Schematic of FANA ASO FLI1 knock-down experiment in OCI-AML3 cells. **H)** The CD34 gene expression level at 3 days post FLI1 knock-down in sg*NT* and sg*STAG2* OCI-AML3 cells. Two independent clones per sg*NT* and sg*STAG2* cells. RNA extracted from CD34 enriched human bone marrow cells are used as control. Bar representing mean ± standard error of mean (s.e.m). **I)** Cell surface CD34 level at 7 days post FLI1 knock-down. Representative FACS plot from two independent experiments. **J)** Venn diagram showing the number of loops detected in sg*NT* and sg*STAG2* OCI-AML3 cells. **K)** The line plot showing the loop gain at CD34 locus in the sg*STAG2* OCI-AML3 cells. The track is overlayed with peaks either unique to FLI1, unique to NPM1c or shared between FLI1 and NPM1c based on CUT&RUN data.

To determine whether these findings could be recapitulated in our isogenic cell line model, we assessed the immunophenotype of our sg*STAG2* AML cells lines to see if there are similar stem cell related cell surface marker changes. We observed significant alterations to the overall immunophenotype of the sg*STAG2* cells with an increase level in CD34 (**Figure 4E, left panel)**, CD123 (**Figure 4E, middle panel)**, and ESAM (**Figure 4E, right panel)** compared to control. To see if FLI1 binding in the sg*STAG2* cells are linked with the increased level of cell surface markers, we examined the chromatin accessibility and FLI1 binding at *CD34* gene. Indeed, there was increased chromatin accessibility and FLI1 binding at a 3’ site with reported DNases hypersensitivity ^46^ that regulates CD34 expression level (**Figure 4F**). Similar trends were observed for FLI1 binding at *ESAM* **(Supplemental Figure 3C)** and *IL3RA* (encoding CD123) promoters (**Supplemental Figure 3D)**.

To test whether FLI1 contributes to CD34 expression, we performed knockdown of FLI1 using 2’-deoxy-2’-fluoro-d-arabinonucleic acid antisense oligonucleotides (FANA ASOs) (**Figure 4G**). FLI depletion resulted in significantly decreased mRNA level of *CD34* (**Figure 4H**) and cell surface protein (**Figure 4I**) after reducing the *FLI1* levels **(Supplemental Figure 3E)** in the sg*STAG2* cells, but not in the control **(Supplemental Figure 3F)**. Finally, we performed Hi-C on the sg*STAG2* and sg*NT* cells to examine the chromatin looping changes. Overall, we observed increased number of loops in the s sg*STAG2* cells (127,991 FitHiC2 loops, 10Kb bin) than sg*NT* cells (85,798 FitHiC2 loops, 10Kb bin) (**Figure 4J**). Among the sg*STAG2* specific loops, we observed a strong association with FLI1 binding at key genes of interest, including *CD34* and *MEN1* (**Figure 4K, Supplemental Figure 3G, Supplemental Table 13)**. These data demonstrate that STAG2 loss in NPM1c AML leads to aberrant FLI1 binding, which drives increased chromatin looping and expression of stem cell program, resulting in a distinct immunophenotypic profile characterized by high CD34 level.

### *Stag2* and *Npm1c* co-mutation results in a fully penetrant and transplantable leukemia model with morphological dysplasia

After characterizing the molecular consequences of STAG2 mutation in human leukemia cell lines, we set out to investigate the effect of co-mutations on leukemogenesis in an *in vivo* setting. To do this, we generated a co-mutant Stag2 and Npm1c conditional activation murine model driven by tamoxifen inducible *Ubc*CreER^T2^ (**Figure 5A**). Upon aging, the *Stag2/Npm1c* co-mutant (referred henceforth as “double”) had much shorter overall survival (median survival time 383.5 days), as well single Npm1c (median survival time 505.5 days), compared to the wild-type and Stag2 single mutant mice (**Figure 5B**). At 52 weeks post mutation activation, the complete blood count (CBC) analysis showed no changes in peripheral white blood cell (WBC) counts (**Figure 5C**), however, the double mutant mice developed macrocytic anemia with decreased red blood cell numbers (**Figure 5D**), decreased hemoglobin (**Figure 5E**) and increased mean corpuscular volume (**Figure 5F**), as well as thrombocytopenia (**Figure 5G**).

**Figure 5:**
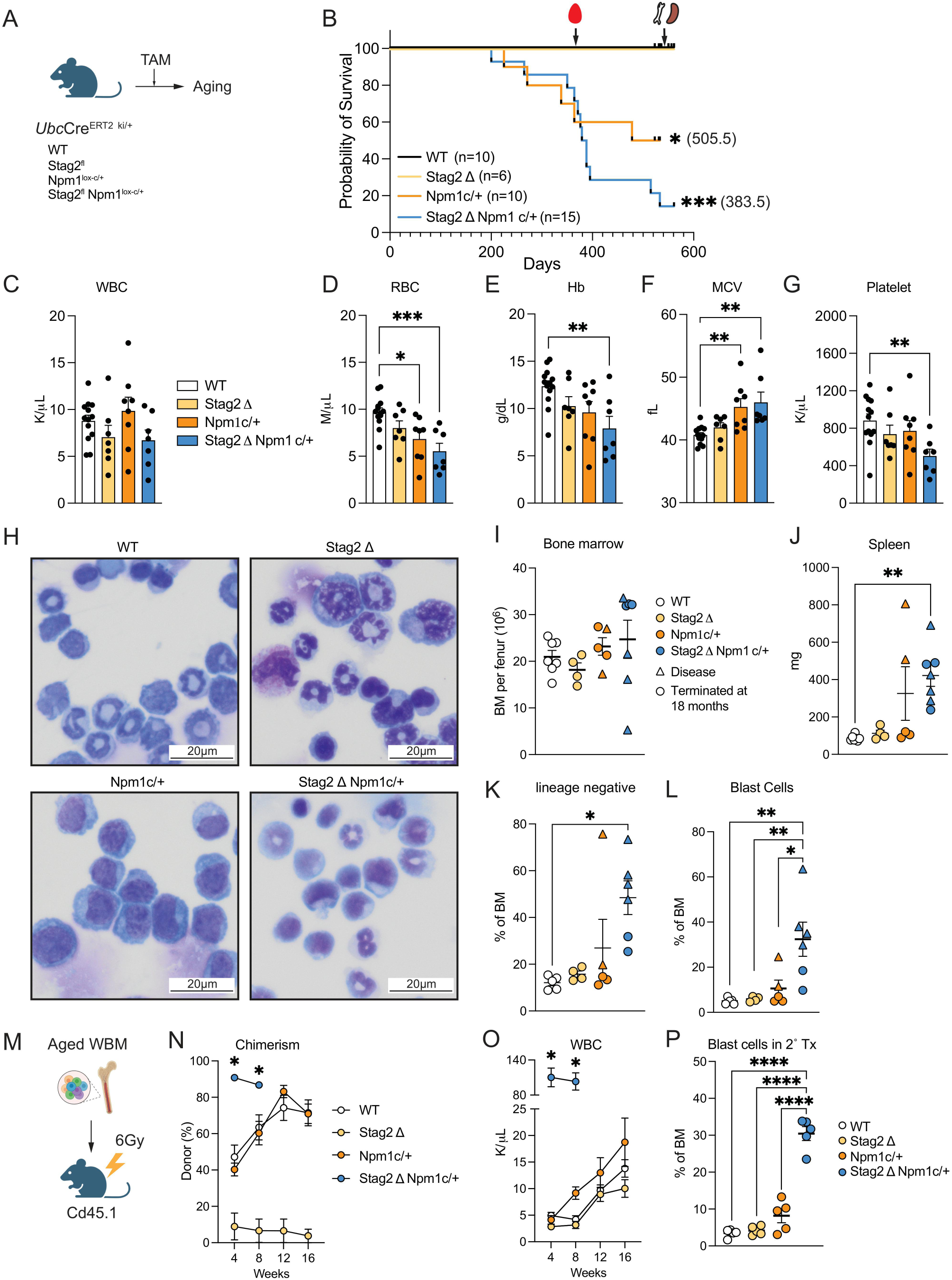
The double Stag2/Npm1c mutation leads to acute leukemia with dysplastic changes. **A)** Schematic of experimental set up. **B)** The Kaplan-Meier survival curves of *Ubc*CreER^T2^ positive WT (control), *Stag2*Δ (single mutant), *Npm1*^c/+^ (single mutant) and *Stag2*Δ *Npm1*^c/+^ (double mutant). *p ≤0.05, ***p ≤0.001, Log-rank test. The median survival time of the Npm1c and Stag2/Npm1c mutant mice are listed in the bracket. **C-G)** Peripheral blood count of control, single mutant and double mutant mice at 1 year post mutation activation: white blood cells (WBC; C), red blood cells (RBC; D), hemoglobin (Hb; E), mean corpuscular volume (MCV; F) and Platelets (G). Data presented as mean±s.e.m. *p ≤0.05, **p ≤0.01, ***p ≤0.001, One-way ANOVA with Tukey’s multiple comparison. **H)** Representative Shandon Kwik-Diff stains of bone marrow cytospin of control, single and double mutant mice at 18 months post mutation activation. Representative micrographs of n=3-5 individual mouse replicates per genotype. All images represented at 40x magnification. Scale bar: 20*_μ_*m. **I)** The bone marrow counts of the aged mice at 18 months post mutation activation of moribund. Data presented as mean±s.e.m. **J)** The spleen weight at the time of the collection. Data presented as mean±s.e.m. **p ≤0.01, One-way ANOVA with Tukey’s multiple comparison. **K)** The percentage of lineage negative population at the time of collection. Data presented as mean±s.e.m. *p ≤0.05, One-way ANOVA with Tukey’s multiple comparison. **L)** The percentage of blast cell population at the time of collection. The blast cells are defined as lineage-FSC^hi^Cd48+Fc*_γ_*R+. Data presented as mean±s.e.m. *p ≤0.05, **p ≤0.01, One-way ANOVA with Tukey’s multiple comparison. **M)** Schematic of aged bone marrow transplantation workflow. **N-O)** The chimerism (N) and WBC count (O) of the transplant recipients of aged bone marrow. n=5 recipients per genotype. Data presented as mean±s.e.m. *p ≤0.05, Two-way ANOVA with Tukey’s multiple comparison. **P)** The percentage of blast cells in the bone marrow of recipients at time of collection. Data presented as mean±s.e.m. ****p ≤0.0001, One-way ANOVA with Tukey’s multiple comparison.

Morphological examination of the bone marrow (BM) cells also indicated presence of myelodysplasia and the appearance of phagocytosing macrophage-like cells in single *Stag2* mutant and the double mutant mice (**Figure 5H, Supplemental Figure 4E)**. When moribund, we observed no changes in overall BM cellularity (**Figure 5I**) while double mutant mice developed significant splenomegaly (**Figure 5J**). Within the BM, the double mutant mice had an expanded lineage negative population (**Figure 5K, Supplemental Figure 4A&C)**, which harbors a distinctive atypical blast cell population (Lin-FSC^hi^Cd48+FcR*_γ_*+) that expanded in the double mutant mice that were present at early time points and markedly expanded upon leukemogenesis (**Figure 5L, Supplemental Figure 4B&D)**.

To confirm the leukemia potential of these cells, we transplanted BM from the moribund mice into sub-lethally irradiated recipients (**Figure 5M**). The double mutant BM recipient rapidly developed acute myeloid leukemia **(Supplemental Figure 4F)**, marked by high engraftment (**Figure 5N**), marked leukocytosis (**Figure 5O**) and myeloid cells **(Supplemental Figure 4G)** in the peripheral blood, as well as splenomegaly **(Supplemental Figure 4H)**. Moreover, the blast cells observed in the primary double mutant mice were significantly expanded in the moribund recipients (**Figure 5P**). Noticeably, double mutant leukemia cells **(Supplemental Figure 4I)**, rather than Npm1c single mutant cells, can propagate in secondary transplant recipients **(Supplemental Figure 4J&K)**. This observation is consistent with prior reports that Npm1c-driven leukemia exhibits limited propagation efficiency in transplantation settings without cooperating genetic lesions^42^. Overall, these findings establish a novel in vivo model of cohesin-mutant leukemia that recapitulates key clinicopathological features observed in patients with cohesin-mutated AML, including myelodysplastic changes and enhanced self-renewal capacity.

### *Stag2* and *Npm1c* co-mutation results in defective hematopoietic stem cell differentiation

Having established the cohesin mutant leukemia model, we set out to characterize the pre-leukemic hematopoietic changes in the HSPC compartment. At 4 weeks post mutation activation, the double mutant mice were thrombocytopenic while no changes observed in the WBC and erythroid panels **(Supplemental Figure 5A-F)**. In the bone marrow, the double mutant mice were hypocellular **(Supplemental Figure 5G)** with expanded hematopoietic stem and progenitor compartment (LSK) **(Supplemental Figure 5H&P)**. Since we observed an expansion of the LSK population, we assessed the cell cycle status and there were no changes between genotypes **(Supplemental Figure 5I)**. Detailed immunophenotypic characterization of the LSK compartment revealed the myelo-erythroid biased MPP2 (LSKFlk2-Cd150+Cd48+) and myeloid biased MPP3 (LSKFlk2-Cd150-Cd48+) were expanded in the double mutant and single Stag2 mutant at the expense of lymphoid biased MPP4 (LSKFlk2+Cd150-) **(Supplemental Figure 5J)**. However, the downstream GMP remains unchanged at this time point of analysis, pointing towards functionally inefficient myeloid commitment **(Supplemental Figure 5K&Q)**. Within the mature populations, we observed increased myeloid cells (Gr1+Cd11b+) at the expense of B (Cd19+B220+) and erythroid (Ter119+Cd71+) population **((Supplemental Figure 5L-N, 5R-T)**. There were no changes in the spleen size for the double mutant mice **(Supplemental Figure 5O)**.

To determine functional changes in the double mutant, we performed colony replating assay on purified LSK and GMP cells, as *Npm1c* mutation was reported to transform not just the hematopoietic stem and progenitor population but also the lineage restricted myeloid progenitor cells ^42^. We found that both the double mutant cells and single *Npm1c* mutant cells had colony replating potential in both LSK **(Supplemental Figure 5U)** and GMP cells **(Supplemental Figure 5V)**. To test the functional changes, we sorted LSK and transplanted cells into lethally irradiated recipients **(Supplemental Figure 6A)**. The double mutant and single *Stag2* LSK cells yielded low peripheral blood chimerism compared to WT **(Supplemental Figure 6B)**. In the donor population, the double mutant cells had a persistently high myeloid percentage **(Supplemental Figure 6C)**. At 16-weeks post transplantation, we compared the chimerism between bone marrow and peripheral blood. We found that despite the low chimerism in the peripheral blood, double mutant *Stag2*/*Npm1c* cells can repopulate the hematopoietic stem and progenitor compartment **(Supplemental Figure 6D)**. When transplanted into secondary recipients **(Supplemental Figure 6E)**, the double mutant LSK cells had poor chimerism **(Supplemental Figure 6F)** and were myeloid biased in the donor portion **(Supplemental Figure 6G)**, consistent with primary transplantation. Overall, the double mutant leads to expansion of the hematopoietic stem and progenitor cells, especially in the myeloid biased MPP compartment in a preleukemia setting. However, the mutant stem and progenitor cells have defective lineage contribution to the blood and myeloid bias, which suggest a differentiation blockade and bone marrow failure characteristic to leukemogenesis ^47^.

**Myeloid biased MPP3 cells are highly dysregulated in the *Stag2/Npm1c* pre-leukemic mice** Our phenotypic and functional analyses identified the LSK compartment as a critical target of Stag2/Npm1c-mediated hematopoietic dysregulation. The LSK population can be further dissected into HSCs or multipotent progenitor cells (MPPs) based on their long-term repopulating ability and lineage output bias. In the *Stag2/Npm1c* mutant there was no changes in HSC quantity while the MPP3 was the most expanded MPP population **(Supplemental Figure 5J)**. Through transplantation studies, we identified a long-term engraftment defect manifest in the blood, as well as myeloid biased output, suggesting likely functional defects in both the stem cell and myeloid bias MPPs. To dissect the changes of the double mutant HSC (LSK Flk2-Cd150+Cd48+) and MPP3 (LSK Flk2-Cd150-Cd48+) population, we performed *in vitro* proliferation assays on purified populations at 4 weeks post mutation activation (**Figure 6A**). Both the double mutant and single *Npm1c* mutant MPP3 cells showed increased proliferation capacity, while *Npm1c* mutant HSCs showed a mild growth advantage compared to all other genotypes (**Figure 6B, Supplemental Figure 7A)**. We then examined differentiation markers and found that the double mutant MPP3 cells had a decreased mature myeloid output compared to WT (**Figure 6C, Supplemental Figure 7B)**. However, we observed no changes in the differentiation capacity of the HSC cultures pointing to an MPP3-specific functional defect **(Supplemental Figure 7C)**.

**Figure 6:**
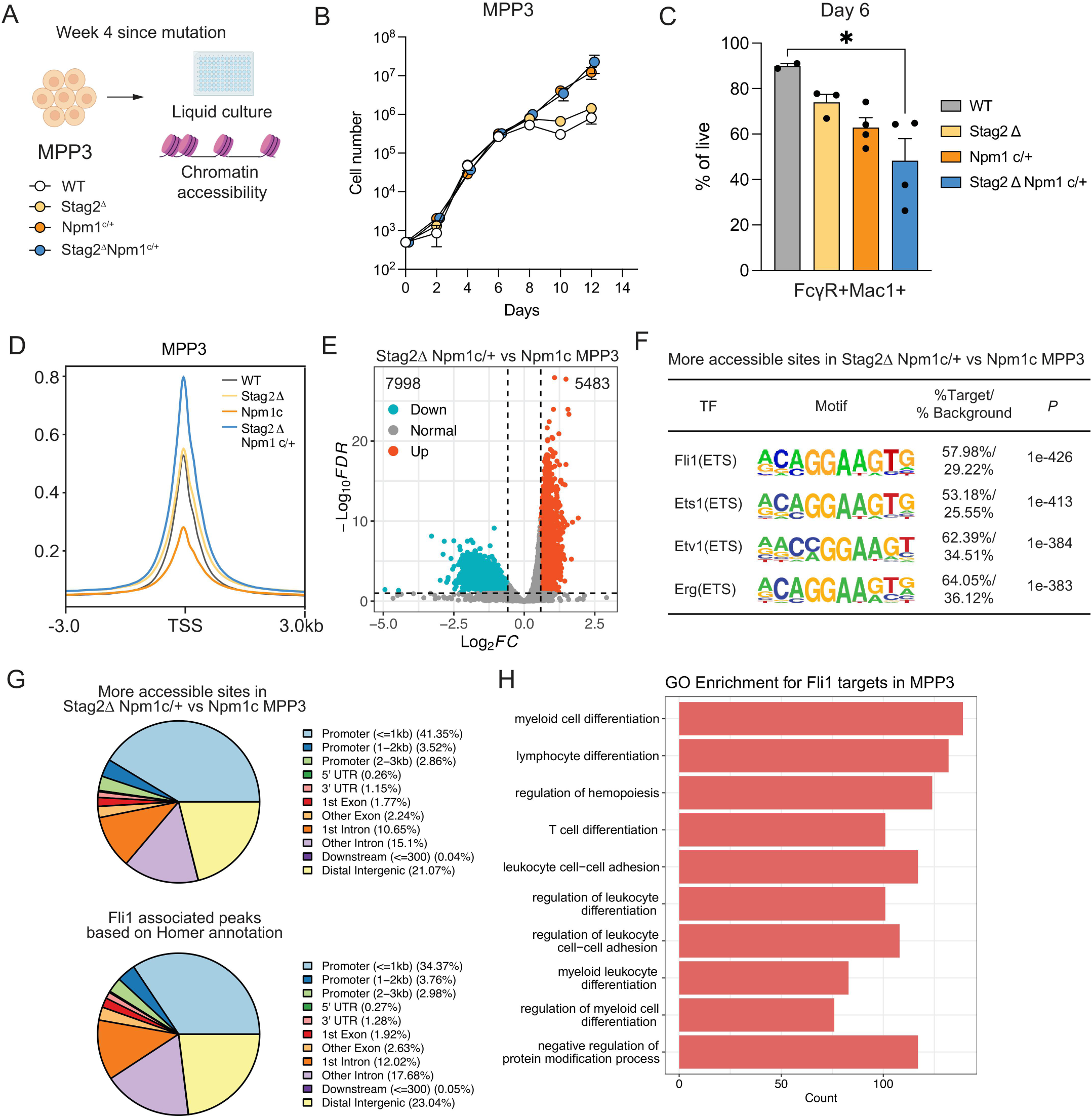
Myeloid biased MPP3 cells are dysregulated in the Stag2Npm1c double mutant mice. **A)** Schematic of experimental set up. **B)** The liquid culture experiment of the MPP3 cells at 4 weeks post mutation activation, n>3 biological replicates per genotype. **C)** The percentage of mature myeloid population (Fc*_γ_*R+Mac1+) cells at day 6 of culture across the genotypes. Bar represents mean±s.e.m. *p ≤0.05, One-way ANOVA with Tukey’s multiple comparison. **D)** The chromatin accessibility of MPP3 cells at transcription start sites (TSS) across the genotypes. N=2-3 biological replicates. Representative plot of two independent datasets. **E)** The differentially accessible sites in the double mutant MPP3 compared to single Npm1c mutant. Cut-off for the differentially accessible peaks are log Fold Change (logFC) > log2(1.5), False Discovery Rate (FDR) < 0.1. **F)** Known homer motif analysis of the more accessible sites in the double mutant MPP3 compared to single Npm1c mutant. For listed TFs, p-value as labelled and q value are 0 (BH-adjusted). **G)** The genomic location of the more accessible peaks in the double mutant compared to single Npm1c mutant (Top panel) and the location where Fli1 motif are identified (Bottom panel). **H)** The GO pathway analysis of the more accessible genes with Fli1 motif in the double mutant cells. Pathways are ranked by adjusted p values.

Given the established link between chromatin structure and cellular identity, we performed bulk ATACseq on the purified HSC and MPP3 cells and found that the double mutation resulted in increased chromatin accessibility in both cell types compared to the WT and single mutant Npm1c (**Figure 6D, Supplemental Figure 7D)**. As the *Stag2/Npm1c* mutation leads to both excessive proliferation as well as a distinct differentiation defect in the MPP3 cells compared to WT, we then compared the chromatin accessibility of the double mutant with single Npm1c as well as with WT MPP3 cells. When compared to WT MPP3, there are fewer accessibility changes (168 up vs 333 down) **(Supplemental Figure 7E)**. However, when compared against single Npm1c mutation, we found that there are 5,483 sites with increased accessibility while 7998 sites are more closed (**Figure 6E**). Homer motif analysis identified that ETS family transcription factor Fli1 is among the top motifs identified in differentially accessible open chromatin (**Figure 6F**). Among the sites that were opened in double mutant vs single Npm1c mutant, the newly accessible MPP3 sites are mostly located around promoter regions (47.73%), followed by Intron (25.75%) and distal intergenic region (21.07%) (**Figure 6G, upper panel)**. As Fli1 is the top motif associated with the more accessible sites, we perform genomic annotation of the Fli1 targeted sites that are more accessible in *Stag2Npm1c* MPP3 and found that these sites are largely located in promoters (**Figure 6G, lower panel)** and around genes involved in differentiation and hematopoiesis regulation (**Figure 6H, Supplemental Table 14)**.

Lastly, we compared the accessibility changes between HSC and MPP3 within the genotype. We found that MPP3s exhibit more accessible sites globally compared to HSC in the WT, while Stag2 mutation leads to a more accessible state in HSC but not in the MPP3 cells **(Supplemental Figure 7F)**, suggesting cell type specific effects of cohesin loss on chromatin architecture.

These findings provide a high-resolution analysis of the functional and epigenomic consequences of Stag2/Npm1c co-mutation in hematopoietic stem and progenitor subpopulations. We identified myeloid-biased MPP3 cells as a key cellular compartment exhibiting enhanced proliferation, impaired differentiation, and extensive chromatin remodeling characterized by increased accessibility at Fli1 target sites. These molecular alterations provide a mechanistic link between aberrant Fli1-mediated transcriptional regulation and the functional hematopoietic dysregulation observed in our pre-leukemic model.

### STAG2 loss enhances dependency on Menin-regulated transcriptional programs

We next evaluated sensitivity to Menin inhibition using Revumenib^48^, a Menin-MLL interaction inhibitor, recently approved for treatment of MLL-rearranged and NPM1c leukemia^49–51^. We reasoned that the altered Menin binding dynamics in the sg*STAG2* could result in differentiation sensitivity towards the Menin inhibitor. We firstly performed dose-response analysis and revealed that sg*STAG2* cells exhibited significantly enhanced sensitivity to Revumenib compared to sg*NT* controls, as evidenced by a lower IC50 value (12 nM vs 36 nM respectively, p<0.0001, **Figure 7A**). After treatment for 7 days, we performed western blotting on the chromatin fraction and observed more Menin binding on the chromatin of sg*STAG2* cells than sg*NT* in the DMSO control. Upon treatment, sg*STAG2* cells had more efficient eviction of chromatin-bound Menin (**Figure 7B**). Taken together, these data demonstrate increased biological responsiveness of sgSTAG2 cells to Revumenib treatment.

**Figure 7:**
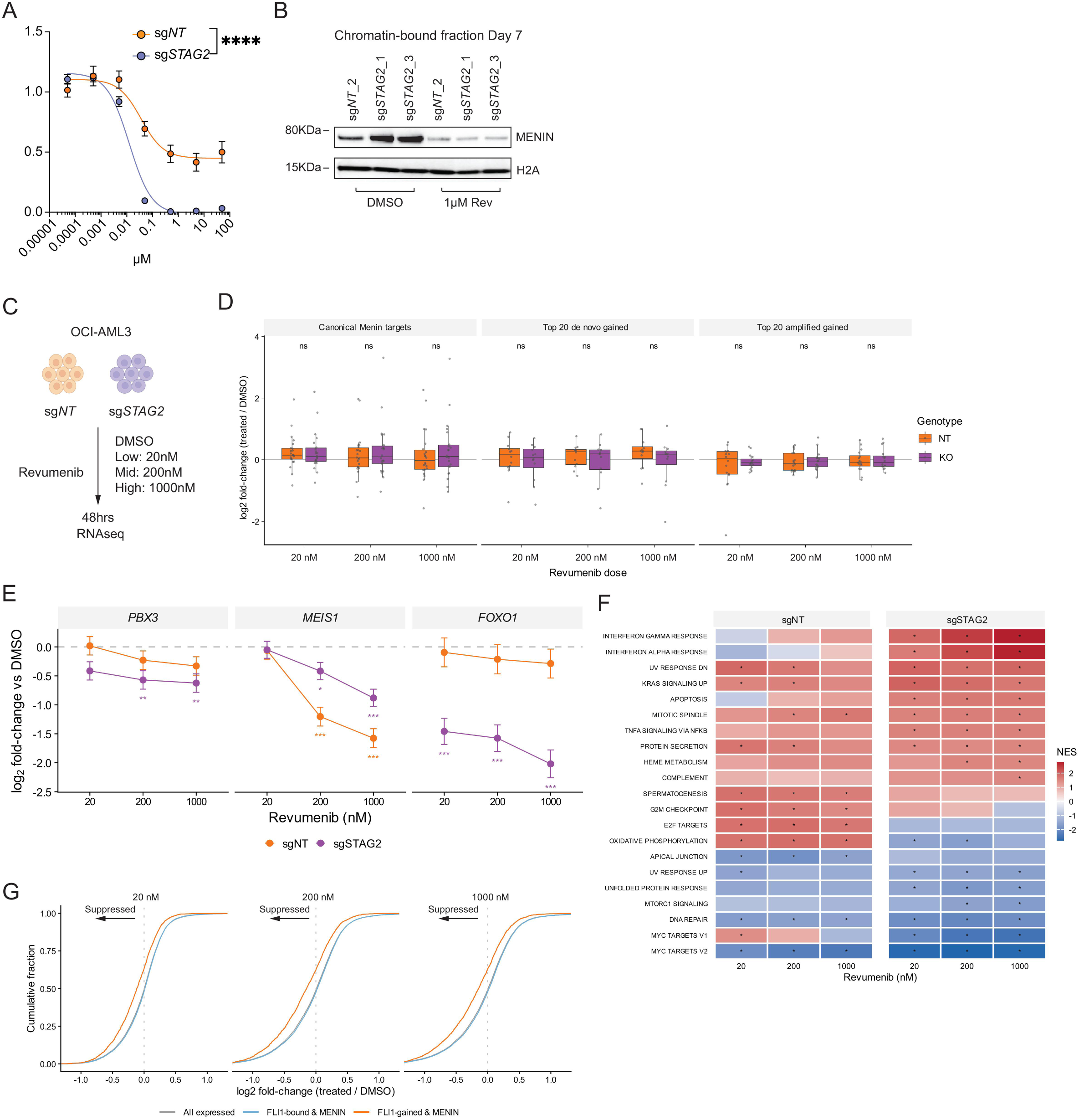
STAG2 mutation alters response to Revumenib in NPM1c leukemia. **A)** Dose-response curves of sg*NT* and sg*STAG2* OCI-AML3 cells to Revumenib at 7 days post treatment. Viabilities are displayed relative to the DMSO control (mean±s.e.m). ****p ≤0.0001, Two-way ANOVA. **B)** Western blot of the chromatin bound fraction collected from sg*NT* or sg*STAG2* OCI-AML3 cells at 7 days post 1μM Revumenib. **C)** Schematic of the low-, mid-and high-dose Revumenib treatment experiment of sg*NT* and sg*STAG2* OCI-AML3 cells. **D)** The transcription response of canonical Menin target genes, the top 20 de novo gained and the top 20 amplified gained genes towards multiple doses of Revumenib. Data is per-gene log2 fold-change against DMSO control. Paired Wilcoxon with Benjamini–Hochberg correction. **E)** The gene expression changes of *PBX3*, *MEIS1* and *FOXO1* gene upon Revumenib treatment in sg*NT* and sg*STAG2*. shown as DESeq2 log₂ fold-change versus DMSO. Error bars represent the standard error of the log₂ fold-change. *p < 0.05, **p < 0.01, ***p < 0.001, two-sided Wald test with Benjamini– Hochberg correction. **F)** Hallmark pathway response to Revumenib in sg*NT* versus sg*STAG2*. Heatmap of GSEA normalized enrichment scores (NES) across Revumenib doses (20, 200, 1000 nM) in sg*NT* and sg*STAG2*. Red, induced by drug (positive NES); blue, suppressed (negative NES). Asterisks mark FDR < 0.05. Shown are pathways reaching FDR < 0.05 at 1000 nM in at least one genotype. **G)** The transcriptome response at low, mid and high dose of Revumenib in *sgSTAG2* OCI-AML3 cells, separated by all genes in grey, FLI1/Menin-bound genes in blue or FLI1-gained and Menin-bound genes in orange.

Our initial analyses revealed that STAG2 loss globally amplifies Menin-dependent signaling, but it is unknown if the amplified signal translates into gene expression changes at baseline or when exposed to Revumenib. To investigate the transcription response of sg*STAG2* cells towards Menin inhibition, we performed low, mid and high dose Revumenib treatment, along with DMSO control, on sg*NT* and sg*STAG2* cells and collected cells for RNAseq at 48 hours post treatment (**Figure 7C**). Based on the Menin binding profile, we quantified the gene expression changes of canonical, top 20 amplified and top 20 de novo Menin bound genes in sg*STAG2* vs sg*NT* across the different conditions. We curated a list of canonical Menin target genes reported from the MLL complex studies^49,52,53^ and excluded genes without detectable Menin binding in the sg*NT* OCI-AML3 cells. Then we selected the top 20 amplified genes, based on Log2FC of Menin signal at sites that overlaps with pre-existing sg*NT* -called peak, as well as the top 20 *de novo* genes, which is a gained site in sg*STAG2* with no peak called in the sg*NT*.

At both baseline and treatment, there are no difference in gene expression among the three categories, while the *de novo* category has the lowest gene expression at baseline (**Figure 7D, Supplemental Figure 8A)**. These data suggest that increased Menin occupancy alone does not explain the enhanced response observed in sg*STAG2* cells. This is further supported by the divergent transcription response observed on the canonical targets. While most genes do not change expression, we found that *PBX3* had a stronger suppression in sg*STAG2*, while *MEIS1* is less responsive compared to sg*NT* (**Figure 7E**). In contrast, the HOX loci have increased gene expression upon treatment in both genotypes **(Supplemental Figure 8B)**.

To explore the molecular basis for increased Revumenib sensitivity, we queried the top 20 genes that are most up-regulated or down-regulated (based on log2 fold-change) upon treatment in sg*NT* and sg*STAG2* OCI-AML3 cells **(Supplemental Figure 8C)**. The pathway analysis on these genes indicates that sgSTAG2 cells have consistent up-regulation of interferon signature, with a suppression of proliferation and MYC programs (**Figure 7F, Supplemental Figure 8D)**. On the gene level, there is a strong downregulation of *FOXO1*, who is among the top 20 de novo Menin-gained genes (**Figure 7E**). Finally, we found a global suppression of FLI1 gained (padj < 0.05, log2FC >1) and Menin bound genes in sg*STAG2* comparing to all expressed, indicating that FLI1-gained/Menin-bound genes are preferentially suppressed following Revumenib treatment. (**Figure 7G**).

Taken together, these findings reveal a mechanistic basis for STAG2-mediated leukemogenesis in NPM1c-mutant AML, where altered chromatin architecture enables ectopic FLI1 binding and expansion of Menin occupancy at regulatory chromatin. The enhanced Menin and FLI1 binding in open chromatin create a vulnerability to pharmacological Menin inhibition, which provides new clues to differential sensitivity observed in the clinic^48,54^.

## Discussion

Hematopoietic differentiation is a tightly regulated, life-long process that sustains regeneration of functional blood cells and is disrupted in leukemia^1,3,4^. The initiation of differentiation in hematopoietic stem cells is controlled by intrinsic molecular mechanisms, including response to receptor signaling^55^, transcriptional activation^11^, and epigenetic priming^6^. In this study, we demonstrate that loss of STAG2 remodels chromatin accessibility to enable ectopic FLI1 binding, promoting Menin enrichment at regulatory chromatin and activation of stemness-associated transcriptional programs in NPM1c-mutant leukemia.

Previous studies have shown that NPM1c establishes oncogenic transcriptional programs through aberrant chromatin binding and dependence on the Menin-MLL complex and have noted enrichment of ETS-family motifs, specifically FLI1 motifs, at NPM1c-bound regions^40,41^. However, the mechanistic basis for this enrichment has not been fully explored. Here we show that STAG2 loss increases chromatin accessibility at FLI1-motifs without altering transcription factor abundance, resulting in expanded FLI1 occupancy across the genome. These newly accessible FLI1-bound regions overlap with NPM1c-and Menin-associated loci and are accompanied by amplification of Menin occupancy across the cistrome, such as *PBX3*, and expansion of Menin binding beyond the canonical HOXA/MEIS1 program. These findings support a model in which STAG2-dependent chromatin remodeling reshapes the Menin-binding landscape in NPM1c AML and leads to an altered transcriptional response towards the Menin inhibition.

While NPM1c mutation is often considered a unifying biomarker for Menin inhibitor eligibility, the clinical application of these agents remains context-dependent, with current approvals largely restricted to the relapsed/refractory setting and ongoing investigation in the frontline setting, particularly in molecularly defined subgroups such as those co-mutated with FLT3-ITD^56^. These observations highlight that NPM1c-mutant AML is not biologically or therapeutically uniform and suggest that additional genomic and epigenomic features—such as STAG2 loss—may refine patient selection and response prediction. In this context, our findings provide a mechanistic framework linking altered chromatin architecture to Menin inhibitor sensitivity, suggesting that STAG2-mediated chromatin remodeling and FLI1–Menin cooperativity may represent a previously unrecognized determinant of response heterogeneity within NPM1c-mutant AML.

Our study also demonstrated that chromatin rewiring is associated with activation of stem cell– like transcriptional programs and an altered immunophenotype characterized by increased *CD34* expression in both cell line models and patient samples. The direct binding of FLI1 at regulatory regions controlling *CD34* and other stemness-associated genes provides a mechanistic link between altered chromatin accessibility and maintenance of a stem cell–like state. These observations are consistent with prior studies demonstrating that cohesin mutations enforce stem cell programs and impair differentiation and extend these findings by identifying a specific transcription factor–mediated mechanism driving this phenotype.

The interaction between chromatin architecture and transcription factor binding illustrated here has broader implications for cancer biology. Oncogenic transcriptional programs are often driven not only by mutated transcription factors or signaling pathways, but also alterations in the chromatin landscape that dictate where these factors bind. In this context, STAG2 functions as a gatekeeper of chromatin accessibility, and its loss reprograms the cistrome of endogenous transcription factors, such as FLI1. Similar mechanisms may underlie other malignancies in which chromatin regulatory mutations co-occur with transcription factor dysregulation. For example, Ewing sarcoma, driven by a FLI1::EWS translocation often acquires STAG2 mutations, which are associated with more aggressive disease^57^. This suggests that chromatin remodeling may broadly amplify oncogenic transcription factor activity across cancer types.

Notably, while STAG2-deficient OCI-AML3 cells demonstrated enhanced responsiveness to Menin inhibition *in vitro*, these studies at current are insufficient to establish STAG2 mutation as a predictive biomarker of clinical response. Future studies will be needed to define the factors governing Menin redistribution in STAG2-deficient leukemia and clinical therapeutic index to Menin inhibition.

Clinically, these findings highlight that NPM1c-mutant AML is not biologically uniform^58^ and that co-occurring mutations such as *STAG2* loss can significantly influence disease phenotype and therapeutic response^30^.The enhanced sensitivity to menin inhibition observed in Stag2/Npm1c co-mutant AML support a model in which chromatin context refines both biological behavior and treatment vulnerability. Given prior studies linking monocytic differentiation states with Venetoclax resistance, the altered differentiation phenotype observed in STAG2-mutant leukemia raises the possibility that cohesin mutations may influence responsiveness to additional therapeutic classes such as the BCL-2 inhibitor Venetoclax^59^. ^59^

Taken together, our findings define a model in which STAG2 loss remodels chromatin accessibility to amplify endogenous transcription factor activity, enabling cooperation with oncogenic drivers such as NPM1c and Menin to promote leukemogenesis. This rewiring of the transcriptional landscape both sustains malignant cell identity and creates specific therapeutic dependencies. More broadly, our findings support a paradigm in which mutations in chromatin architectural regulators drive cancer through context-dependent reprogramming of transcription factor cistromes, revealing new opportunities for precision therapeutic interventions.

## Resource availability Lead contact

Requests for further information and resources should be directed to and will be fulfilled by the lead contact, Aaron D. Viny

## Materials availability

All unique/stable reagents generated in this study are available from the lead contact without restriction.

## Data and code availability

Processed sequencing data is made available via NCBI Gene-Expression Omnibus (GEO) at GSE292654,GSE292660. Other data generated in this study are available within the article and its supplementary files, or on request from the corresponding author. The NPM1c Cut&Run data analyzed in this study were obtained from GSE197387. The Mx1 Stag2 LSK and GMP RNAseq and ATACseq data were obtained from GSE134583. Codes used for data analysis are available upon request.

## Acknowledgements

We are grateful to members of the Viny lab and Columbia Stem Cell Initiative (CSCI) for their discussion of this work. This work was funded by NIH/NCI grants R37 CA286857 (A.D. Viny) and a Scholar award from the American Society of Hematology (A.D. Viny). A.D. Viny is a Damon Runyon-Doris Duke Clinical Investigator supported by the Damon Runyon Cancer Research Foundation (CI-120-22) and a Clinician Scientist Development Award from the Doris Duke Charitable Foundation. J.J.X. is supported by a postdoctoral fellowship by the Walker Family and the Edward P. Evans Foundation Columbia MDS Center. V.Scoca is a recipient of a postdoctoral fellowship from the American Italian Cancer Foundation and Blood Cancer United (LLS) career development fellow. V.S.Sudunagunta was supported by an HONORS award from the American Society of Hematology and a Physician Scientist Support Foundation award from the American Society for Clinical Investigation. We are grateful for the support from the Abel Family Foundation and the Lewis and Spear-Wein Families. This research was funded in part through the NIH/NCI Cancer Center Support Grant (P30CA013696) and supported by the Clinical and Translational Science Awards (CTSA) grant (UL1TR001873) from the National Center for Advancing Translational Sciences (NCATS). We thank the Sulzburger Genome Center/Genomics and High Throughput Drug Screening at the Herbert Irving Comprehensive Cancer Center (HICCC), Flow Cytometry Core at the CSCI of Columbia University Irving Medical Center. Schematics included in the figures were generated on BioRender. We acknowledge the Addgene for the plasmid distribution.

## Author’s Contributions

**J.J. Xu**: Conceptualization, data curation, software, formal analysis, funding acquisition, validation, investigation, visualization, methodology, writing–original draft, project administration, writing–review and editing. **V. Scoca**: Data curation, formal analysis, investigation, visualization, validation, methodology, writing–review and editing. **Y. Chen**: Data curation, software, formal analysis, investigation, visualization, validation. **Y.A. Zhan**: Investigation, visualization. **A. Fisher**: Investigation, visualization. **E. Udoh**: Investigation, visualization. **S. Fernando**: Investigation, visualization, validation, methodology. **B. Alija**: Investigation. **J. Pantazi**: Investigation. **V. Sudunagunta**: Investigation. **E. Stewart**: Investigation. **A.M.D. Galang**: Investigation. **M. Williams**: Investigation, visualization, methodology. **G. Bhagat**: Investigation, visualization, methodology. **C. Gebhard**: Data curation, formal analysis, methodology, supervision. **V. Visconte**: Data curation, formal analysis, methodology. **S. Ondrejka**: Data curation, formal analysis, visualization, methodology. **R. Delwel**: Data curation, formal analysis, methodology, supervision. **M. Hu**: Data curation, software, formal analysis, investigation, visualization, methodology. **R.P. Koche**: Data curation, formal analysis, investigation, methodology. **A.D. Viny**: Conceptualization, data curation, formal analysis, supervision, funding acquisition, investigation, visualization, methodology, writing–original draft, project administration, writing–review and editing.

## Methods

### Mice

All animal experiments were conducted at the Columbia University Irving Medical Center (CUIMC) in accordance with Institutional Animal Care and Use Committee protocols approved at CUIMC and ethics regulation. *Mx1*-Cre ^60^, *Ubc*CreER^T2^ ^61^, *Stag2*^fl^ ^27^ and *Npm1*^flox-c/+^ ^62^ mice were previously described and were bred in house to generate the *Ubc*CreER^T2^ *Stag2*^fl^ *Npm1*^flox-c/+^ cross. All mice were aged in house. 6-12 weeks old WT B6.SJL-Ptprca Pepcb/BoyJ (Ptprca, Jax Laboratory, #:002014) or WT C57BL6-CD45.1/2 mice were used for transplantation experiments. WT C57BL6-CD45.1/2 mice were bred in house by crossing WT C57BL/6-CD45.2 (B6) and Ptprca mice. Both male and female animals were used in discriminately in all experiments. Animal facilities were maintained at 22±2 °C and 50±10% relative humidity on a 12-12h light-dark cycle. Mice were euthanized by CO_2_ asphyxiation followed by cervical dislocation.

### *In vivo* experiments and transplantation assays

*Ubc*CreER^T2^ positive wild type, *Stag2*^fl^ and *Npm1*^flox-c/+^, and *Stag2*^fl^ *Npm1*^flox-c/+^ mice were tamoxifen treated via oral gavage (100mg/kg, dissolved in corn oil) for 1-2 doses when aged 6-8 weeks old. *Mx1*-Cre *Stag2*^fl^ or *Stag2*^fl^ mice were treated with Polyinosinic:polycytidylic acid (pIpC, GE Healthcare) as described previously ^27^. For timepoint analysis, peripheral blood was isolated by submandibular bleeds into EDTA-containing tube (Becton-Dickinson) for analyses on a Genesis (Oxford Science) hematology system per manufacturer’s instruction. Bone marrow (BM) cellularity was determined using a ViCell automated cell counter (Beckman-Coulter). Spleen were weighted on a chemical balance scale.

For transplantation experiments, CD45.1 or CD45.1/2 recipient mice were exposed to 6Gy (sublethal, 1 dose) or 10.5Gy (lethal, split dose, 3hrs apart) irradiation delivered using an X-ray irradiator (MultiRad225, Precision X-Ray Irradiation), and purified LSK, or unfractionated BM cells were delivered by retro-orbital injections. For LSK transplantation in to lethally irradiated recipients and secondary transplantation of LSK cells, 10,000 LSK were delivered together with 200,000 unfractionated CD45.1 supporter BM cells. For competitive BM transplantation, 1×10^6^ mutant BM were mixed with 1×10^6^ CD45.1 competitor BM cells and delivered into recipients. For sublethal transplantation, 2×10^6^ aged BM cells were delivered into recipients. For secondary sublethal transplantation of leukemic BM, mice received 1×10^6^ mutant BM were mixed with 1×10^6^ CD45.1 competitor BM cells. Lethally irradiated recipient mice were administered with water containing polymyxin (1MU/L, P4932-5MU, Sigma) and neomycin trisulfate salt hydrate (1.2mg/mL, N1876, Sigma) for 4 weeks following transplantation.

### Flow cytometry, cell sorting, and western blotting

For immunophenotyping analysis, peripheral blood was ACK (150mM NH_4_CI, 10mM KHCO_3_ and 0.1mM Na_2_EDTA) lysed to remove red blood cells (RBCs) and washed once with Phosphate buffered saline solution (PBS). BM cells were obtained by flushing both femurs in staining medium composed of PBS containing 2% heat-inactivated Fetal Bovine Serum (FBS, #MT35016CV, Corning). BM RBCs were removed by lysis with ACK and washed once with staining medium. For cell sorting experiment, BM cells were obtained by crushing hindlimb and pelvic bones in the staining medium and removed RBCs. BM cells were then pre-enriched for lineage-cells using EasySep™ Mouse Hematopoietic Progenitor Cell Isolation Kit (Stem Cell Technology). For hematopoietic stem and progenitor cell analysis and sorting, unfractionated or lineage-enriched BM cells were then incubated with Cd4 PE-Cy5 (eBioscience, 1:800), Cd11b PE-Cy5 (eBioscience, 1:800), Gr1 PE-Cy5 (eBioscience, 1:800), Cd8 PE-Cy5 (eBioscience, 1:400), B220 PE-Cy5 (eBioscience, 1:400), Ter119 PE-Cy5 (eBioscience, 1:200), Cd3 PE-Cy5 (eBioscience, 1:100), Sca1 PE-Cy7 (Biolegend, 1:400), c-Kit APC-Cy7 (Biolegend, 1:400), Cd150 PE (Biolegend, 1:100), Cd48 Af700 (Biolegend, 1:100) for immunophenotyping or Cd48 Af647 (Biolegend, 1:100) for sorting, Cd34 FITC (eBioscience, 1:200), Fc*_γ_*R BV711 (Biolegend, 1:200) and Flk2 biotin (Biolegend, 1:100) followed by SA BV421 (Biolegend, 1:400). For peripheral blood immunophenotyping, ACK-lysed cells were incubated with B220 PE-Cy7 (Biolegend, 1:200), Mac1 Af700 (Biolegend, 1:1000), Gr1 PE (Biolegend, 1:300), Cd3 PerCP-Cy5.5 (Biolegend, 1:200), Cd4 APC-Cy7 (Biolegend, 1:300), Cd8a FITC (Biolegend, 1:200) and ckit APC (Biolegend, 1:200). For mature BM cell analysis, unfractionated BM cells were incubated with B220 PE-Cy7, Mac1 Af700, Gr1 PE, Cd19 APC-Cy7 (Biolegend, 1:400), Cd4 FITC (Biolegend, 1:200), Cd8a FITC and ckit APC. For peripheral blood and bone marrow mature population chimerism analysis, cells were incubated with B220 PE-Cy7, Mac1 Af700, Gr1 PB (Biolegend, 1:1000), Cd3 PerCP-Cy5.5, Cd4 FITC, Cd8a FITC, c-Kit APC-Cy7, Cd45.2 APC (Biolegend, 1:100), Cd45.1 PE (Biolegend, 1:100). For hematopoietic stem and progenitor chimerism analysis and sorting, BM cells were incubated with Cd4 PE-Cy5, Cd11b PE-Cy5, Gr1 PE-Cy5, Cd8 PE-Cy5, B220 PE-Cy5, Ter119 PE-Cy5, Cd3 PE-Cy5, Sca1 PE-Cy7, c-Kit APC-Cy7, Cd150 BV605 (Biolegend, 1:100), Cd48 Af700, Cd34 FITC, Fc *γ* R BV711, Flk2 biotin followed by SA BV421. For *in vitro* differentiation assays, cultured cells were stained with c-Kit APC, Sca1 PB (Biolegend, 1:400), Cd150 PE, Cd48 Af700, Mac-1 APC-Cy7 (Biolegend, 1:400) and FcγR BV711 (Biolegend; 1:400). For OCI-AML3 cells surface marker analysis, cultured cells were stained with anti-human CD11b PE (555388, BD,1:200) or ESAM PE (MA5-46656, Invitrogen, 1:200), CD34 Af700 (343526, Biolegend, 1:200) and CD123 PerCP-Cy5.5 (306016, Biolegend, 1:200). Before analyses, stained cells were resuspended in staining medium containing 1 μg ml^−1^ propidium iodide or DAPI (4’,6-diamidino-2-phenylindole) for dead cell exclusion. Cell analyses were performed on the Becton Dickinson (BD) FACS ARIA II SORP, Novocyte Quanteon and Novocyte Penteon. Cell sorting was performed on FACS ARIA II SORP or SONY MA900. Data collection was performed using FACSDiva (v.9), NovoExpress (v2.2.) or SONY MA900 software and analysis was performed using FLowJo (v10). For clinical flow cytometry, all cases were immunophenotyped using either 6-color FACSCanto II (dates 2018-2019) or 12-color Lyric (dates 2020-2023) flow cytometry instruments with FACSuite software (Becton Dickinson, San Jose, CA) and analyzed with FCS Express (DeNovo Software). Blast populations were identified by CD45 and side scatter characteristics and the following antibodies were used to characterize blasts (all obtained from Becton Dickinson): CD45-APC/Cy7, CD34-APC, CD13-PE/Cy7, HLADR-PerCP/Cy5.5, CD33-PE, and CD65-FITC.

For western blotting, protein lysates were extracted from harvested OCI-AML3 pellets using RIPA buffer (78840, ThermoFisher Scientific) containing a cOmplete^TM^, EDTA free protease inhibitor cocktail (Roche). For subcellular fractionation, protein lysates were prepared from 10^7^ cells using Subcellular Protein Fractionation Kit for Cultured Cells (ThermoFisher Scientific) as per manufacturer’s instruction. Protein quantification was performed using the Pierce^TM^ Bradford protein assay (23200, ThermoFisher Scientific) and analyzed on a Luminex 200 analyzer as per manufacturer’s instruction. For co-immunoprecipitation, 100ug of chromatin fraction were pre-cleared with 1ug IgG negative control antibody for Cut&Run (13-0042, Epicypher) overnight. The pre-cleared lysates were then incubated with 1ug of rabbit anti-Menin (A300-105A, Bethyl) or IgG overnight. Then proceed with immunoprecipitation kit based on manufactures instructions (ab206996, Abcam). Proteins were then separated by NuPAGE 4-12% Bis-Tris Gel and transferred to a nitrocellulose membrane. After transfer, the presence of protein was checked via Ponceau S (ThermoFisher Scientific) and the membrane was blocked in TBST with 5% Bovine serum albumin (BSA, ThermoFisher Scientific). The following primary antibodies were used: rabbit anti-GAPDH (D16H11, Cell signaling), rabbit anti-SA2 (A302-580A, Bethyl), rabbit anti-Histone H2A (D603A, Cell Signaling), rabbit polyclonal anti-FLI1 (ab153909, Abcam), rabbit anti-Menin. After overnight incubation and washing in 1x TBST, the blots were incubated with secondary anti-rabbit HRP (7074S, Cell Signaling) or Mouse Anti-Rabbit IgG (Light-Chain Specific) (D4W3E, Cell Signaling) Monoclonal Antibody (HRP Conjugate) for co-IP for 1hr, following by washing and developed with ProSignal^TM^ Pico ECL reagent (Genesee Scientific) or SuperSignal™ West Femto Maximum Sensitivity Substrate (ThermoFisher Scientific). Images were obtained using the iBright Chemi Blot (Invitrogen).

### *In vitro* primary murine cell culture and colony forming assays

All cultures were performed at 37°C in 5% CO_2_ water jacket incubator (ThermoFisher). For myeloid differentiation assays, 500 HSC or MPP3 cells were directly sorted into 96 well round bottom plate containing Iscove’s modified Dulbecco’s medium with GlutaMAX^TM^ (Invitrogen), supplemented with 5% FBS, 1% penicillin (50U ml^-1^) and streptomycin (50*_μ_* g ml^-1^), and the following cytokines (all from PeproTech): recombined mouse stem cell factor (SCF; 25ng ml^-1^), recombined mouse thrombopoietin (TPO; 25 ng ml^−1^), recombined mouse Flt3L (25 ng ml^−1^), recombined mouse IL-3 (10 ng ml^−1^)), recombined mouse IL-11 (25 ng ml^−1^), recombined mouse granulocyte–macrophage colony-stimulating factor (GM-CSF; 10 ng ml^−1^) and recombined human erythropoietin (EPO; 4 U ml–1). Cells were analyzed by flow cytometry for cell counts using the absolute count function on Novocyte per manufacturer’s instruction and cell surface marker for different culture periods. For colony forming assays, 1,000 LSK or 10,000 GMP cells were seeded in 1mL MethoCult M3434 (Stem Cell Technologies) with no additional supplemental cytokines in duplicates on 6 well plates and scored on day 7. For replating, cells were harvested and pooled and re-seeded at 1,000 LSK or 10,000 GMP cells/well in 1mL MethoCult M3434 in duplicate.

### Plasmids and lentiviral vector production

LentiCas9-Blast (#52962) and LentiGuide-Puro (#52963) plasmids were purchased from Addgene. STAG2 and Non-targeting (NT) sgRNA sequences ^63^ were cloned into the transfer vector according to published protocol ^64^. LentiCas9-Blast and the lentiGuide-Puro lentiviral vectors were produced by transient transfection of HEK293T cells through calcium chloride coprecipitation in 10 cm dish plates. Co-transfection was performed with 13 µg of transfer vector, 10 µg of psPAX2 (#12260, Addgene), and 3 µg of pCMV-VSV-G (#8454, Addgene) plasmid. After the collection of the supernatant 48 h post-transfection, lentiviral particles were concentrated through Lenti-X concentrator (#631232, Takara) and stored at −80°C. Lentiviral vectors were titrated by qPCR of WPRE sequence in HEK293T cells at 3 days post-transduction.

### CRISPR editing, shRNA knock-down and culture of human leukemia cell line

All cultures were performed at 37°C in 5% CO_2_ water jacket incubator (ThermoFisher). OCI-AML3 leukemic cell line is cultured in MEM alpha media (ThermoFisher) supplemented with 20% FBS and 1X Penicillin-Streptomycin solution (100 U/ml and 100 ug, respectively). A stock of Cas9-positive OCI-AML3 cells was generated through transduction (MOI 0.3) of the cells with LentiCas9-Blast vector and selection at 24h post transduction with 5 µg/mL of Blasticidin for 14 days. CRISPR knock out clones were induced from a second transduction of these cells with STAG2 and Non-targeting (NT) lentiGuide-Puro vectors (MOI 0.3) and selection in both presence of Blasticidin and Puromycin (0.5 µg/mL) for 10 days. Both STAG2 and NT transduced cells were single cell-sorted in 96-well plate until colony growth for screening of protein knock out through intracellular flow cytometry.

The shRNA knock-down of the STAG2 in human CD34+ core blood cells and ATACseq were described previously ^65^. For knock-down of FLI1, 2’-deoxy-2’-fluoro-d-arabinonucleic acid antisense oligonucleotides (FANA ASOs) were synthesized by AUM BioTech, LLC (Philadelphia, PA) using the following target sequence: CGTCATGTTCTGGTTTGAGAT ^66^. On the day of transduction, 1nmole of ASOs or scramble control were added to 0.5 ×10^6^ sg*NT* or sg*STAG2* OCI-AML3 cells in 1mL. The *FLI1* gene expression level was assessed on day 3 post transduction by qPCR and cell surface marker via flow cytometry at day 7.

### *In vitro* drug assay

To determine the response of sg*NT* and sg*STAG2* OCI-AML3 cells towards Menin inhibitor, 10^5^ cells from two independent clones were plated in a 96-well plate in triplicate in 100µL of media. Revumenib (HY-136175, MedChem Express) was added the cells using a D300e Digital Dispenser (Tecan) in the following concentrations (50µM, 10µM, 2µM, 0.4µM, 80nM, 16nM and 3.2nM) or DMSO only. After 7 days, viability was determined using Cell Counting Kit 8 colorimetric assay (HY-K0301, MedChem Express). Viability ratios were calculated as ratios of treatment absorbance over vehicle absorbance and curve fitting was done using Prism (GraphPad) inhibitor vs response model. For measuring cell count and cell surface marker changes towards Revumenib, 10^6^ cells from two independent clones were cultured in 2mL of media containing DMSO only or 2.5 µM Revumenib in a 12 well plate. Cell counting was performed using ViCell every 3 days and cell surface marker were measured vis flow cytometry at day 9 post drug treatment. After counting the cells, 10^6^ cells were resuspended with fresh DMSO or drug at respective concentration and cultured in a 12 well plate.

### Assessment of cell cycle, intracellular STAG2 and BrdU assay

For cell cycle analysis of murine hematopoietic stem and progenitor cells, unfractionated BM cells were incubated with the same FACS antibody panel as hematopoietic stem and progenitor analysis. Then cells were fixed and permeabilized with Invitrogen™ eBioscience™ Foxp3 / transcription factor staining buffer set as per manufacturer’s instructions. Cells were then blocked with purified rat anti-mouse Cd16/32 (93, Biolegend) for 15 mins before staining with Ki-67 FITC (B56, BD) overnight. The next day, cells were washed twice and resuspended in PBS for analysis.

For cell cycle analysis of OCI-AML3 clones, 10^6^ cells were cultured for 1 hour with 10 µM BrdU. After harvesting and fixation, DNase treatment and staining was performed according to BrdU staining kit (8817-6600, eBioscience) protocol and finally resuspended in DAPI. For intracellular STAG2 analysis, cells are washed in PBS and first stained with surface antibodies for OCI-AML3 analysis. Then, cells were incubated with Live/Dead Fixable Dye Near IR (780) (#L34993, Thermo Scientific) for 20 minutes and then fixed with 2% paraformaldehyde (PFA) for 15 minutes. Then, intracellular staining is performed with Anti-STAG2 (ab201451, Abcam, 1:200) in a solution of PBS, 2% BSA and 0.2 % Triton-X for 30 minutes, followed by one wash and secondary Donkey anti-rabbit Af405 (A48258, Thermo Scientific, 1:1000) staining in same buffer. Cells were then washed twice and resuspended in PBS for analysis.

### Histology

To prepare cytopsin, 10^5^ BM cells were spun at 800g for 7 mins using Cytospin 4 cytocentrifuge (ThermoFisher) onto charged slides. Cytospins were fixed with 100% methanol for 10 mins and dried at room temperature for at least an hour. The slides were stained with Shandon Kwik-Diff stains (Epredia, 99-907-00) according to manufacturer’s protocol. Sternum was fixed in 4% paraformaldehyde for over 24hrs. The subsequent sternum tissue processing, section and H&E staining were performed by HIstowiz. Images were acquired using Olympus BX41 inverted microscope at 40X objective magnification.

### RNA sequencing and data analysis

10^6^ cultured sg*NT* or sg*STAG2* OCI-AML3 cells were washed in PBS and lysed in TRIzol (Thermo Scientific). The total RNA was extracted with Zymo Direct-zol RNA Microprep Kit (R2062, Zymo Research) and quantified using RNA ScreenTape on Agilent 4150 Tapestation. The cDNA conversion and library prep were performed using SMART-Seq mRNA LP kit (634769, Takara) and the Unique Dual Index Kit (634752, Takara). For input, 9ng of total RNA were used to generate cDNA. The cDNA was quantified using D5000 ScreenTape on Agilent 4150 Tapestation and 4ng of cDNA was used for the library prep with 14 cycles of PCR amplification. The RNAseq libraries were sequenced on NovoSeq X at >30M read depth.

Raw FASTQ files were processed using the nf-core/rnaseq pipeline (v3.17.0) via Nextflow ^67,68^, with Docker containers ensuring a fully reproducible computational environment. Quality control included adapter trimming and removal of low-quality bases using Trim Galore (v0.6.10), followed by quality assessment with FastQC (v0.12.1). The results were aggregated and visualized using MultiQC (v1.25.1). Reads were aligned to the UCSC mm10 mouse reference genome for LSK/GMP cells and hg38 for OCI-AML3 cells using STAR (v2.6.1d)^69^, guided by corresponding the GTF annotation file. In parallel, Salmon (v1.10.3)^70^ was employed for transcript-level quantification, generating both raw and length-scaled read counts. For differential expression analysis, the length-scaled count matrix from Salmon served as input to DESeq2 ^71^ (v1.28.0). Normalized expression values were estimated using TPM (Transcripts Per Million), and differentially expressed genes (DEGs) were identified based on an adjusted p-value threshold (FDR < 0.05) and a fold-change (FC) cutoff (|log2FC| > log2(1.5)). Additionally, a full ranked gene list (by log2FC) was tested for pathway-level enrichment using Gene Set Enrichment Analysis (GSEA)^72^, incorporating gene sets from msigdbr (category “C2”) and a permutation-based significance assessment.

### ATAC sequencing and data analysis

Sorted HSC or MPP3 cells were washed once in 1mL of ATAC resuspension buffer (RSB, 10mM Tris-HCI pH7.4, 10mM NaCI, 3mM MgCI_2_) and then resuspended in 50µL of ice-cold ATAC Lysis buffer (0.1% NP40, 0.1% Tween-20, 0.01% Digitonin in RSB) and permeabilized for 3mins on ice. The nuclei pellets were then washed once in 1mL ice cold ATAC wash buffer (0.1% Tween-20 in RSB). The nuclei pellets were then resuspended in 50 µL of transposition mix (10mM Tris-HCI pH7.4, 5mM MgCI_2_, 10% Dimethyl Formamide, 100mM Transposase, 0.01% Digitonin, 0.1% Tween-20, 33% PBS and 10% molecular water) and incubated for 30 mins at 37°C, 1000rpm for transposition reaction. Post transposition, cleaved DNA were cleaned using Zymo DNA clean and concentrator kit as per manufacturer’s reaction and eluted in 25µL. Then, purified DNA were amplified in NEBNext 2x master mix and indexes using the following PCR program: 5 mins at 72°C, 30s at 98°C, followed by 14 cycles of 10s at 98°C, 30s at 63°C and 60s at 72°C. Following amplification, the product was purified using Ampure XP beads (Beckman Coulter) at 0.45x ratio first and then 1.17x ratio. Library quantity and size distribution were checked on D1000 or D5000 tape on Agilent 4150 Tapestation (Agilent Techologies). The ATACseq libraries were sequenced on NovoSeq X at >30M read depth.

ATAC-seq data were processed using the nf-core/atacseq pipeline (v2.1.2) in Nextflow. Quality control steps included adapter trimming and quality filtering with Trim Galore, followed by alignment of paired-end reads to the hg38 or mm10 reference genome using Bowtie2 (v2.4.4)^73^. Read coverage was computed with BEDTools (v2.30.0)^74^, normalized to Counts Per Million (CPM), and converted to BigWig format using bedGraphToBigWig for visualization. MACS2 (v2.2.7.1)^75^was employed for narrow peak calling, and read quantification was performed with featureCounts. Differential accessibility analysis was conducted using DESeq2, with significant differentially accessible regions identified based on an FDR < 0.05 and |log2FC| > log2(1.5). To assess chromatin accessibility patterns, global signal distribution, and read coverage, deepTools (v3.5.3)^76^ was used to generate heatmaps and coverage profiles around key genomic features (e.g., transcription start sites). Additionally, HOMER (v5.1)^77^ was employed for motif discovery and enrichment analysis, scanning peak regions for enriched DNA motifs and annotate motif associated genes. The GO analysis was performed using Cluster Profiler (v4.10.1) in R.

### CUT&RUN and data analysis

Cleavage Under Targets and Release Using Nuclease (CUT&RUN) were performed using CUTANA^TM^ ChIC/CUT&RUN Kit and CUT&RUN Library Prep kit (EpiCypher) as per manufacturer’s instructions using 500,000 cells per target and 5ng of CUT&RUN DNA for library prep. The following antibody were used for the CUT&RUN: rabbit polyclonal anti-FLI1 (ab153909, Abcam), rabbit anti-Menin antibody (A300-105A, Bethyl), H3K27ac Antibody for CUT &RUN (13-0059, EpiCypher), H3K4me3 Antibody for CUT &RUN (13-0060, EpiCypher) and IgG negative control antibody for Cut&Run (13-0042, Epicypher). Library quantity and size distribution were checked on D5000 tape on Agilent 4150 Tapestation (Agilent Technologies). The CUT&RUN libraries were sequenced on NovoSeq X at >30M read depth.

CUT&RUN data were processed using the nf-core/cutandrun pipeline (v3.2.2) in Nextflow. Raw reads underwent adapter trimming and quality filtering, followed by alignment to the hg38 reference genome using Bowtie2. A blacklist file was applied to remove known sequencing artifacts and nonspecific binding regions. Peak calling was performed using both MACS2(v2.2.7.1) and SEACR (v1.3)^78^ to identify narrow and broad peaks. Read counts were normalized using the CPM method, with control samples incorporated for background correction. For differential peak analysis, DESeq2 was used, and global signal distribution, read coverage, and motif enrichments was further visualized through deepTools and HOMER, following a workflow similar to the ATAC-seq analysis.

### Hi-C and data analysis

Both mouse LSK and GMP cells and CRISPR edited OCI-AML3 cells were isolated for Hi-C library preparation using the Arima Hi-C kit as previously described using low cell input modifications ^27^. Briefly, cells were cross-linked using 1% methanol-free formaldehyde for 10 minutes and quenched in 125mM glycine. Cross-linked cells were lysed, and chromatin was restriction enzyme digested using restriction enzymes that digest chromatin at ^GATC and G^ANTC, where N can be any of the 4 genomic bases (Arima Genomics, San Diego, CA). Digested chromatin was reverse crosslinked using NaCl and eluted in 20uL 2X Shearing buffer (Covaris, Woburn, MA) and fragmented to 350 base pair fragments using a Covaris LE220Rsc sonicator (Covaris, Woburn, MA). Sheared genomic material was biotinylated and enriched using streptavidin beads. Genomic libraries we prepared to streptavidin-bound DNA using Arima protocol modifications for Accel-NGS 2S DNA plus library kit (IDT, Coralville, IA). After end repair and ligation, libraries were quantified using the KAPA library quantification kit (Roche, Indianapolis, IN) and PCR amplified for the number of cycles required to generate >4nM per library. Hi-C libraries were sequenced for on an Illumina NovoSeq X at 500M read depth and raw sequencing data in the Fastq format were obtained.

To analyze Hi-C data we first used Chromap ^79^ to map raw fastq files to the mm10 and hg38 reference genomes, for mouse and human Hi-C data, respectively. We only kept uniquely mapped reads and removed PCR duplicates. Next, we binned mapped reads into 50Kb resolution Hi-C contact matrices and applied the HiCrep ^80^ method to evaluate the reproducibility between biological replicates. After confirming the high reproducibility between biological replicates, we combined biological replicates together and used pooled data for all the downstream analysis.

We first calculated the contact probability curve, i.e., the relationship between 1D genomic distance and average Hi-C contact frequency, as described in the previous study ^32^. Next, we used the principal component analysis method to define A/B compartments at 50Kb bin resolution^32^. We further identified the boundary of topologically associating domains at 50Kb bin resolution, using the insulation score method ^81^. We also identified the frequently interacting regions (FIREs) at 50Kb bin resolution, as described in our previous work ^82^. Lastly, we used HiCCUPS ^31^ to identify 10Kb resolution chromatin loops, and FitHiC2 ^83^ to identify 10Kb resolution long-range enhancer-promoter interactions.

### Quantitative real-time PCR

Total RNA was extracted from cells lysed in the TRZol solution using the Zymo Direct-zol RNA Microprep Kit (R2062, Zymo Research) per manufacturers’ protocols. Complementary DNA was then reverse transcribed using the Verso cDNA Synthesis Kit (ThermoFisher). hFLI1 (Hs00956709) and hCD34 (Hs02576480) was evaluated by quantitative reverse-transcription (qRT) PCR using Taqman probes on the CFX Opus Real-Time PCR systems (BioRad). The gene expression levels were normalized against hGAPDH (Hs02786624).

### Statistical analysis

All experiments were repeated as indicated; *n* or the number of data points indicate the number of independent biological replicates. Mice used for tamoxifen treatment, aging post mutation activation and transplantation were assigned to experimental groups based on genotype and randomized with respect to sex. Data were processed using Microsoft Excel (v.16). Statistical significance was evaluated using GraphPad Prism (v.10) and R (v 4.3.3). Data distribution was assumed to be normal but not formally tested and outliers were not excluded unless indicated.

## Supplemental Figure Legends

**Supplemental Figure 1:**
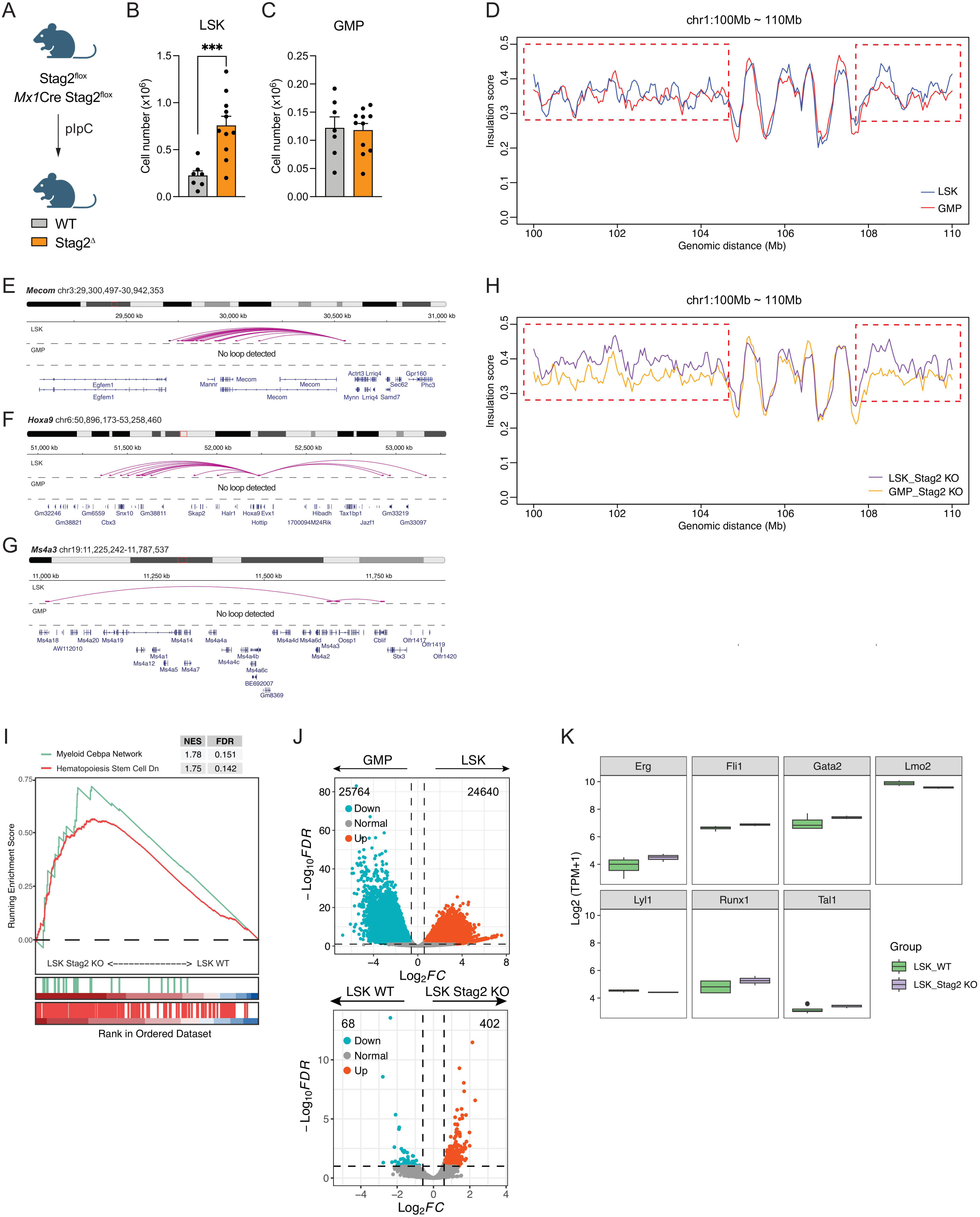
Dynamic chromatin remodeling between hematopoietic stem and progenitor cells and myeloid progenitors is not altered by Stag2 loss. Related to Figure 1 & 2. **A)** Schematic of experimental set up. **B-C)** The number of LSK (**B**) and GMP (**C**) in the bone marrow at 6 weeks post Stag2 deletion. Data presented as means.e.m, ***p 0.001, Unpaired student t-test. **D)** The insulation scoring of 10Mb region of chromosome 1 of WT LSK and GMP cells. The red dashed square highlights the region with high contact frequency in the LSK cells. **E-G)** Lineplot of Mecom loops **(E)**, Hoxa9 **(F)**, Ms4a3 **(G)** in LSK and GMP cells. **H)** The insulation scoring of 10Mb region of chromosome 1 of Stag2Δ LSK and GMP cells. The red dashed square highlights the region with high contact frequency in the LSK cells. **I)** The GSEA analysis of WT LSK and Stag2 LSK with enrichment scoring of stem cell and myeloid Cebpa network. **J)** The chromatin accessibility comparing WT LSK and WT GMP cells (upper panel) and comparing WT LSK and Stag2 LSK cells (lower panel) (n=3 biological replicates). **K)** The expression of heptad transcription (TF) factors (*Fli1, Erg, Gata2, Runx1, Tal1, Lyl1, Lmo2*) in WT and Stag2 LSK cells.

**Supplemental Figure 2:**
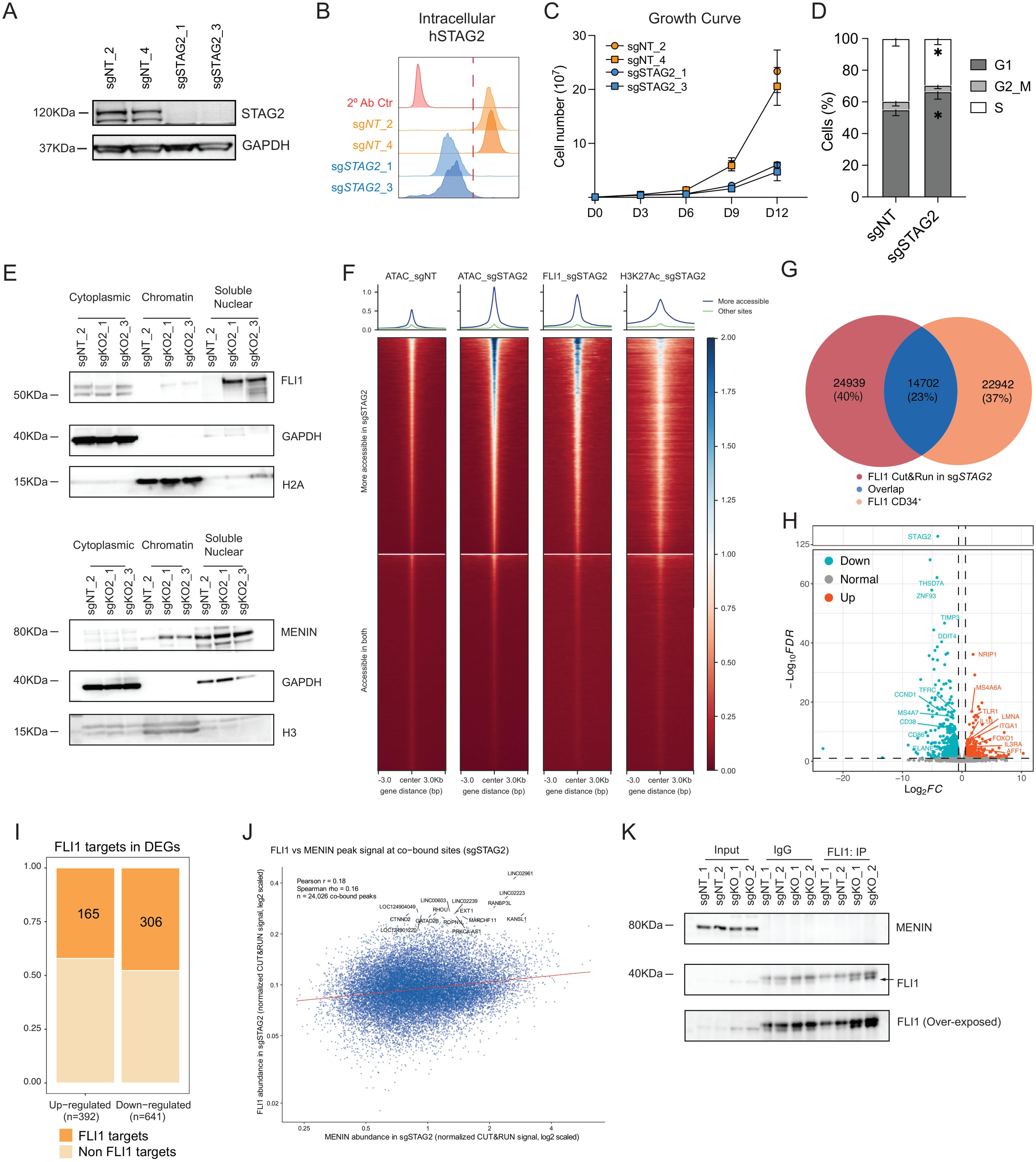
Characterization of the STAG2 KO OCI-AML3 cell lines. Related to Figure 2&3. **A)** Western blotting of STAG2 and GAPDH control in non-targeting (sg*NT*) control and STAG2 KO (sg*STAG2*) OCI-AML3 cell lines. **B)** Intracellular flow cytometry analysis of sg*NT* and sg*STAG2* OCI-AML3 cells. **C)** The proliferation of sg*NT* and sg*STAG2* OCI-AML3 cells. Data presented as means.e.m. **D)** The cell cycle profile of sg*NT* and sg*STAG2* OCI-AML cells. Data presented as means.e.m. *p 0.05, Unpaired student t-test. **E)** Western blotting of FLI1 (upper panel) or Menin (lower panel) in the subcellular fraction of sgNT and sgSTAG2 OCI-AML3 cells. **F)** Heatmap of chromatin accessibility of sgNT and sgSTAG2, FLI1 and H3K27Ac Cut&Run in sgSTAG2, grouped by more accessible region of sgSTAG2 or other peaks. **G)** Intersection of FLI1 binding between sgSTAG2 OCI-AML3 cells with published CD34+ dataset. **H)** The differentially expressed genes in sg*STAG2* comparing to sg*NT*. **I)** FLI1 target genes defined by the consensus FLI1 peak binding at the promoter sites (<1Kb) of differentially expressed genes. Cut-off for the differentially expressed genes are log Fold Change (log2FC) > log2(1.5), False Discovery Rate (FDR) < 0.1. **J)** The FLI1 and Menin signal at co-bound peaks in sgSTAG2 OCI-AML3 cells. Red line represents the linear fit. **K)** Co-Immunoprecipitation of FLI1 in sgNT and sgSTAG2 chromatin fraction, blotted for Menin and FLI1.

**Supplemental Figure 3:**
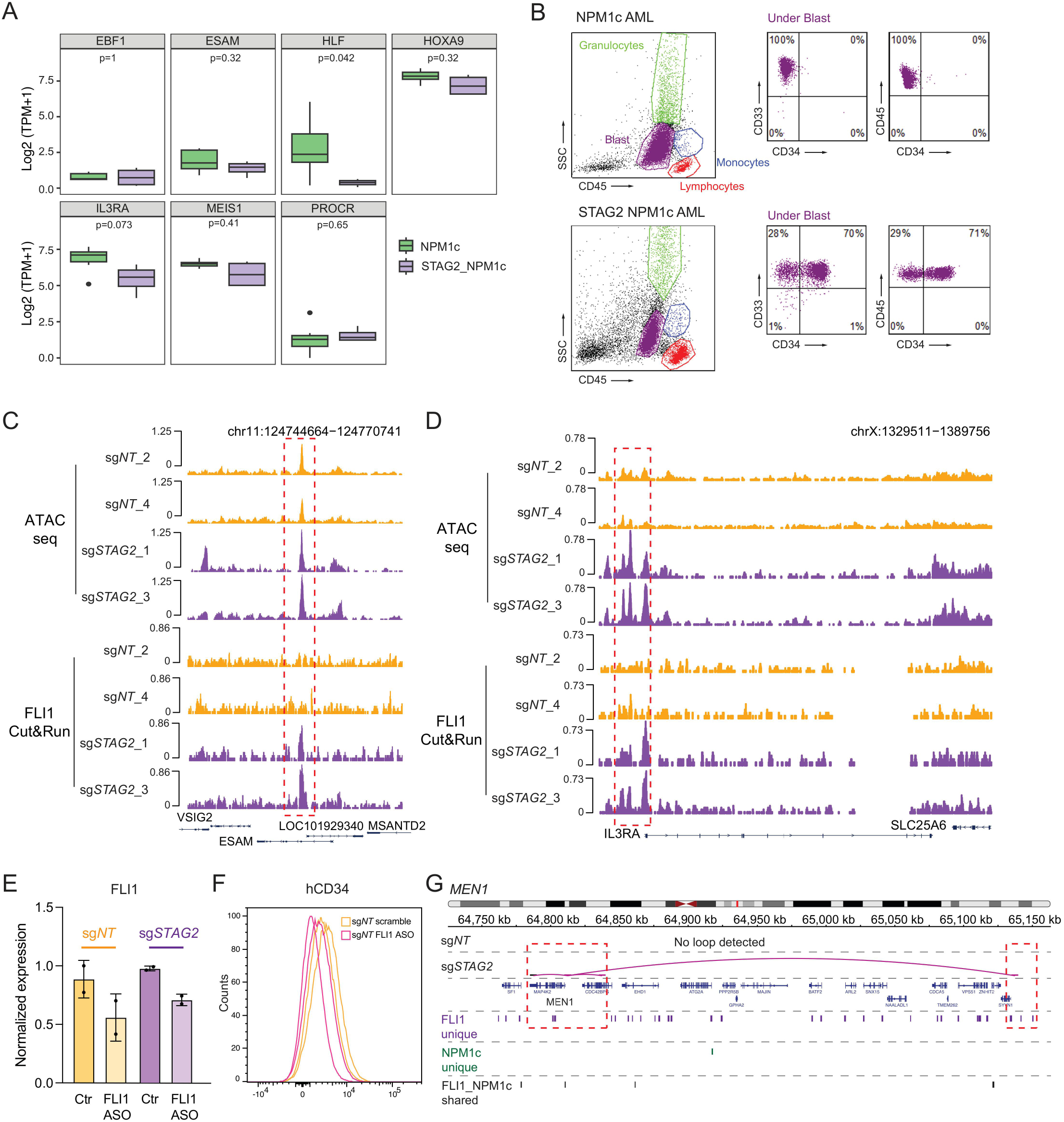
Additional characterization of the FLI1 localization in the STAG2 KO OCI-AML3 cells and associated cell surface markers. Related to figure 4. **A)** The gene expression level of *EBF1, ESAM, HLF, HOXA9, IL3RA, MEIS1, PROCR* in NPM1c mutant (n=6) and STAG2 NPM1c mutant (n=4) AML patients. *p* value displayed is calculated by Wilcoxon signed rank test. **B)** Representative FACS plot showing the gating strategy of NPM1c and STAG2 NPM1c mutant AML. The gated population include blasts (Purple), granulocytes (Green), monocytes (Blue), lymphocytes (Red). The CD34 levels are gated under blast cells. **C-D)** The IGV track view of ATACseq and FLI1 Cut&Run binding in sg*NT* and sg*STAG2* OCI-AML3 cells on *ESAM* (C) and *IL3RA* (gene encoding CD123) locus (D). The red box highlights the area with FLI1 binding in sgSTAG2 cells. **E)** The real-time qPCR level of FLI1 in control (Scramble) and FLI1 ASO of sg*NT* and sg*STAG2* at 3 days post FLI1 knock-down. Data presented as means.e.m. **F)** Representative FACS plot showing cell surface CD34 level at day 7 post FLI1 knock-down in sg*NT* OCI-AML3 cells. **G)** The line plot showing the loop gain at *MEN1* locus in the sg*STAG2* OCI-AML3 cells. The track is overlayed with peaks either unique to FLI1, unique to NPM1c or shared between FLI1 and NPM1c based on Cut&Run data. The red box highlights the area with more interactions and the adjacent peaks in sgSTAG2 cells.

**Supplemental Figure 4:**
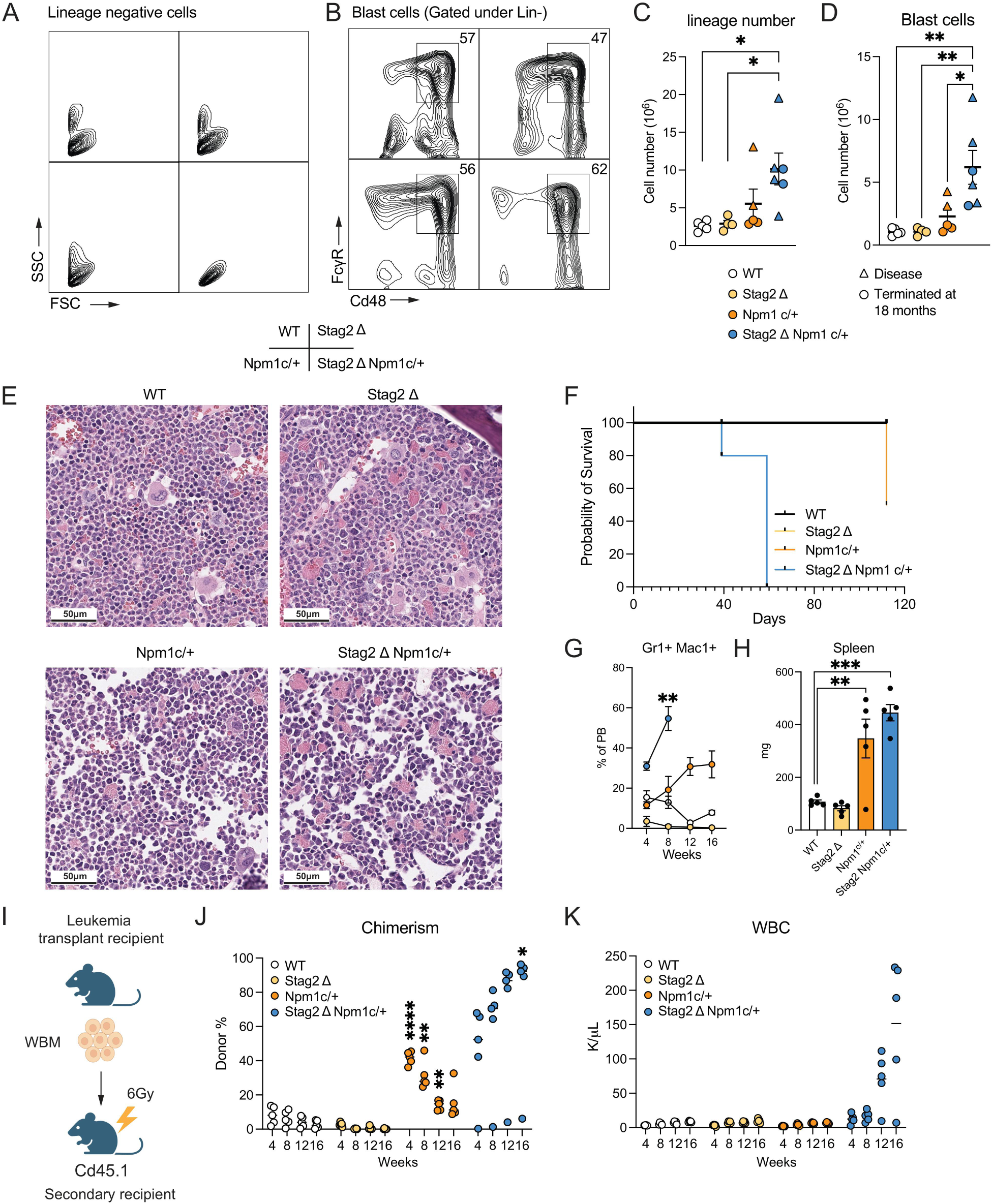
The double Stag2Npm1c mutation leads to AML with dysplastic change. Related to figure 5. **A-B)** Representative FACS plot of FSC and SSC profile of the lineage negative cells (A) and blast population (B) of UbcCre positive WT, Stag2, Npm1c and Stag2Npm1c mutant at 18 months post mutation activation. **C)** The number of lineage negative cells in the bone marrow of 18 months aged or moribund mice. Data presented as means.e.m. *p 0.05. One-way ANOVA with multiple comparison. **D)** The number of blast cells (Lineage-FSC^hi^Cd48+FcR+) in the bone marrow of 18 months aged or moribund mice. Data presented as means.e.m. *p 0.05, **p 0.01, One-way ANOVA with multiple comparison. **E)** Representative hematoxylin and eosin (H&E) of sternum bone marrow of 18 months aged UbcCre positive WT, Stag2, Npm1c and Stag2Npm1c. Images taken at 40x magnification. Scale bar: 50m. **F)** Kaplan-Meier survival analysis of the transplantation recipients of aged control or mutant bone marrow. n=5 recipients per genotype. **G)** The percentage of granulocytes (Gr1+Mac1) cells in the donor (Cd45.2+) fraction in the transplant recipients. Data presented as means.e.m. **p 0.01, One-way ANOVA with multiple comparison. **H)** The spleen weight of transplant recipients at time of sacrifice. Data presented as means.e.m. **p 0.01, ***p 0.001 One-way ANOVA with multiple comparison. **I)** Schematic of experimental workflow of the secondary transplantation of leukemic bone marrow. **J-K)** The chimerism (J) and white blood cell (WBC, K) of the secondary transplant recipients. Data are presented as individual points with mean. **p 0.01, ****p 0.0001, Two-way ANOVA with multiple comparison. Significance displayed is calculated against wild type (WT).

**Supplemental Figure 5:**
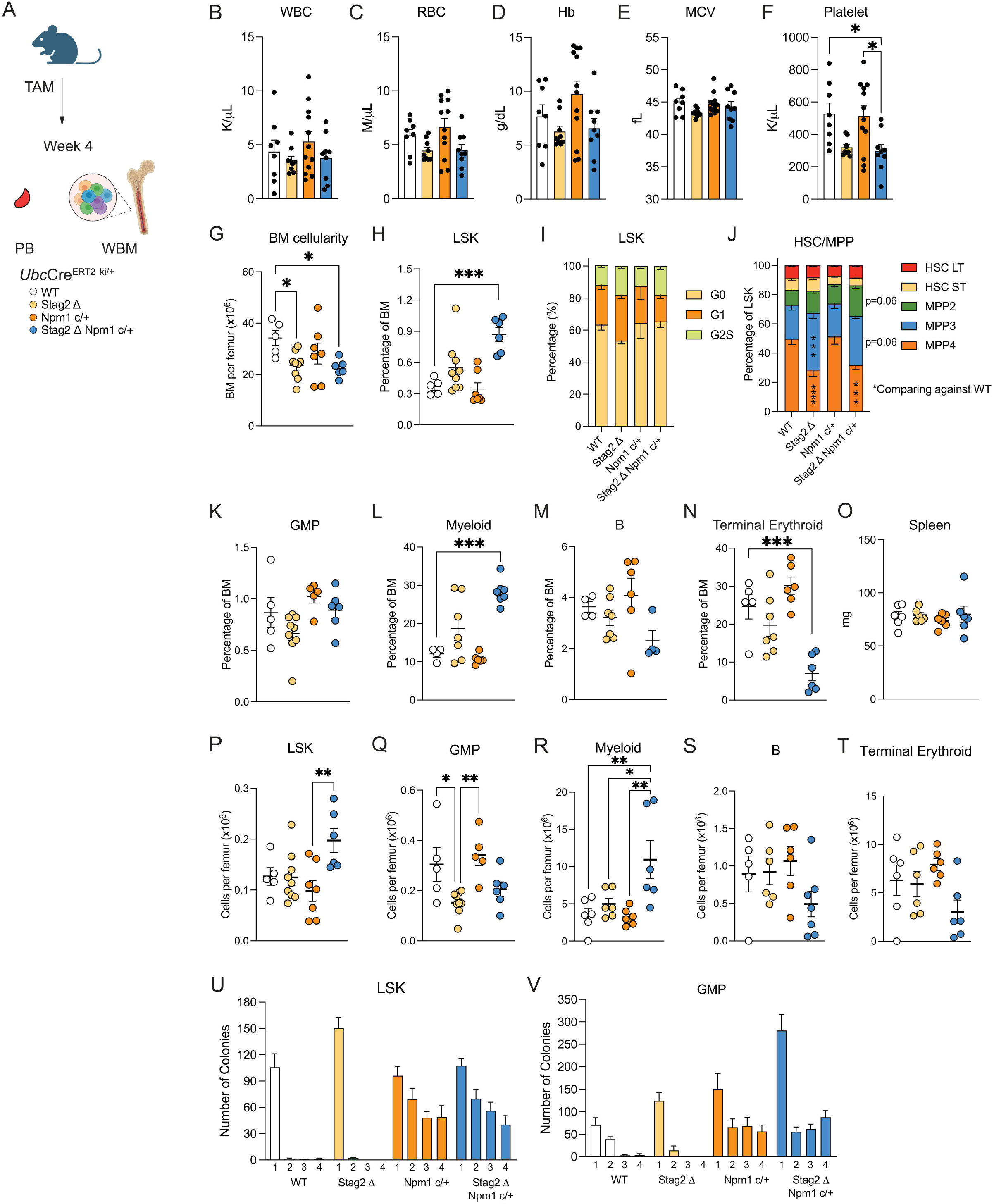
Stag2Npm1c mutation induces expansion of hematopoietic compartment with increased self-renewal capacity. **A)** Schematic of experimental set up. **B-F)** Peripheral blood count of control, single mutant and double mutant mice at four weeks post mutation activation: white blood cells (WBC; B), red blood cells (RBC; C), hemoglobin (Hb; D), mean corpuscular volume (MCV; E) and Platelets (F). **G)** The bone marrow counts of *Ubc*-CreER^T2^ positive WT (control), *Stag2* (single mutant), *Npm1*^c/+^ (single mutant) and *Stag2 Npm1*^c/+^ (double mutant) mice at 4 weeks post mutation activation. **H)** The percentage of LSK (Lineage-Sca1+cKit+) cells in the bone marrow. **I)** The cell cycle analysis of the LSK compartment of the mutant mice at 4 weeks post mutation activation. **J)** The percentage of long-term hematopoietic stem cells (HSCLT, LSKFlk2-Cd150+Cd48-), short-term hematopoietic stem cells (HSCST, LSKFlk2-Cd150-Cd48-), multipotent progenitor 2 (MPP2, LSKFlk2-Cd150+Cd48+), multipotent progenitor 3 (MPP3, LSKFlk2-Cd150-Cd48+), and multipotent progenitor 4 (MPP4, LSKFlk2+Cd150-) cells under the LSK population, Two-way ANOVA with Tukey’s multiple comparison. **K)** The percentage of granulocytic-macrophage progenitor (GMP, Lineage-cKit+Sca1-Cd34+FcR+) in the bone marrow. Data presented as means.e.m. **L-N)** The percentage of mature myeloid (Gr1+Mac1+, L), B lymphoid (B220+Cd19+, M) and terminal erythroid (Ter119+Cd71+, N) population in the bone marrow. **O)** The spleen weight of mutant mice at 4 weeks post mutation activation. **P)** The absolute number of LSK cells in the bone marrow. **Q)** The absolute number of GMP in the bone marrow. **R-T)** The absolute number of mature myeloid (R), B lymphoid (S) and terminal erythroid (T) population in the bone marrow. **U-V)** Serial replating assay of mutant LSK (P) or GMP (Q) compartment at 4 weeks post mutation and scored at day 7 after each replating (sample plated in duplicate, means.e.m). Data presented as means.e.m in this figure, *p 0.05, **p 0.01, ***p 0.001, ****p 0.0001, One-way ANOVA with Tukey’s multiple comparison unless specified.

**Supplemental Figure 6:**
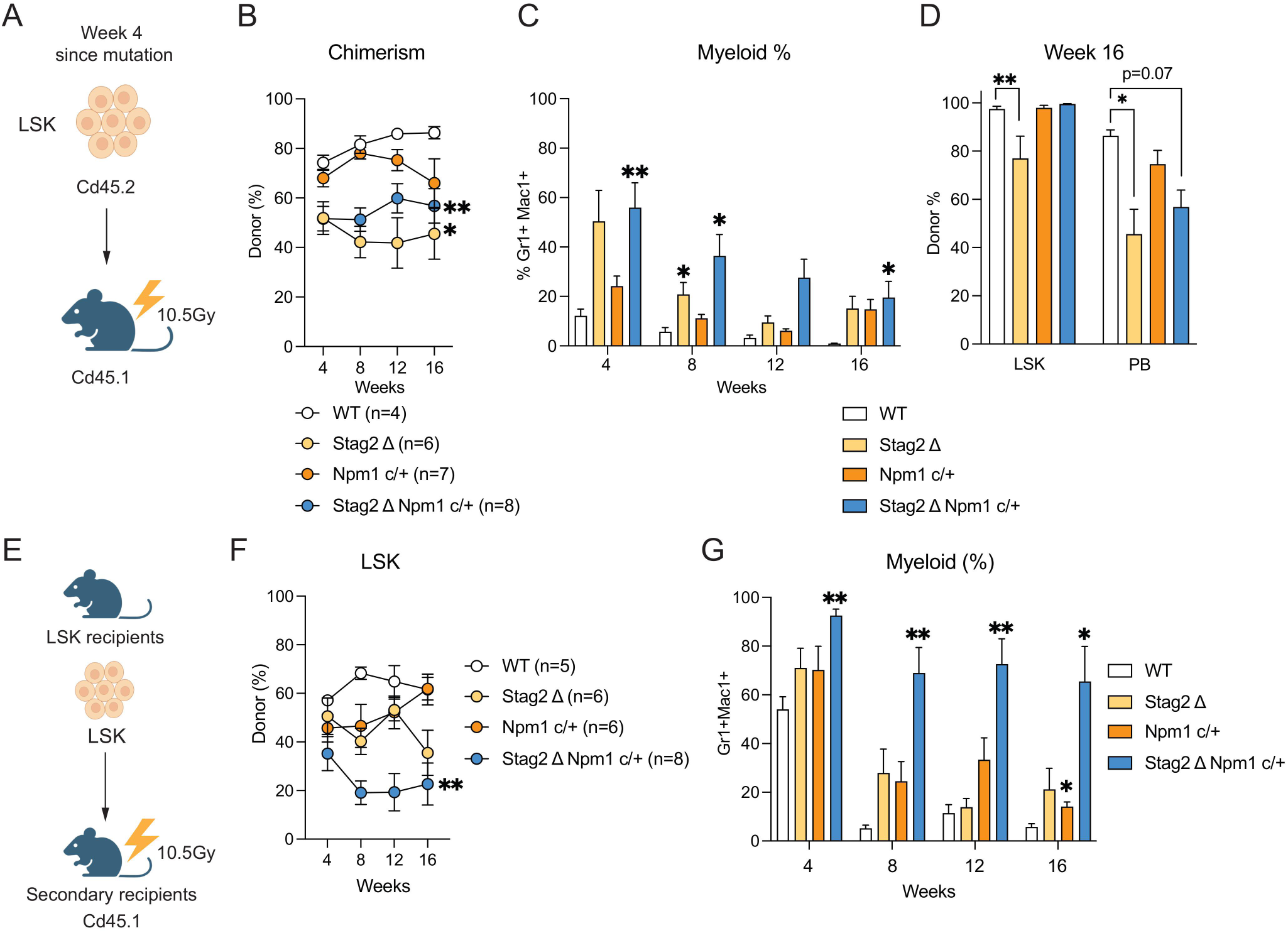
Stag2Npm1c mutant cells are myeloid biased and have poor engraftment capacity in the peripheral blood. **A)** Schematic of LSK transplantation experiment. **B)** The chimerism of transplant recipients of LSK cells in the peripheral blood over 16 weeks post transplantation. Data presented as means.e.m, *p 0.05, **p 0.01, Mixed model ANOVA with multiple comparison. **C)** The percentage of myeloid (Gr1+Mac1+) population in the donor fraction (Cd45.2+) of the recipients. Bar represents means.e.m, *p 0.05, **p 0.01, Mixed model ANOVA with multiple comparison. **D)** The percentage of donor chimerism (Cd45.2%) in the LSK fraction and the peripheral blood (PB). Bar represents means.e.m, *p 0.05, **p 0.01, One-way ANOVA with Tukey’s multiple comparison. **E)** Schematic of secondary transplantation of the LSK cells. **F)** The chimerism of secondary LSK transplantation in the peripheral blood. Data presented as means.e.m. **p 0.01, Two-way ANOVA with multiple comparison. **G)** The percentage of myeloid (Gr1+Mac1+) cells in the donor (Cd45.2+) fraction of recipients. Data presented as means.e.m. *p 0.05, **p 0.01, Two-way ANOVA with multiple comparison.

**Supplemental Figure 7:**
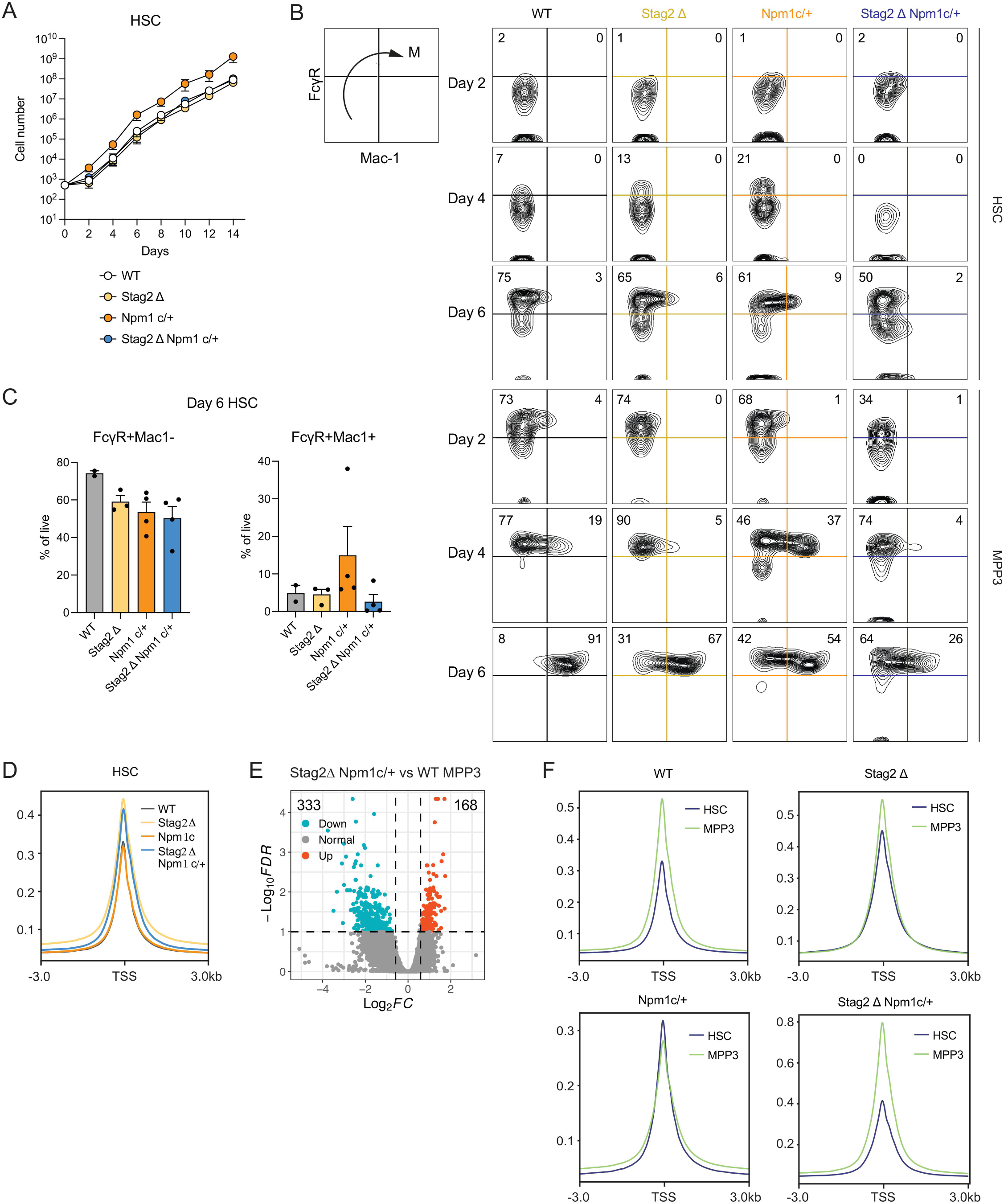
Myeloid biased MPP3 cells are dysregulated in the Stag2Npm1c double mutant mice. Related to figure 6. **A)** Cell counts of the *Ubc*Cre positive WT, Stag2, Npm1c and Stag2Npm1c mutant HSC in liquid culture. Data presented as means.e.m., n>3 per genotype. **B)** Representative FACS plots showing the kinetics of FcR+ and Mac1 acquisition in the HSC and MPP3 population of each genotype. **C)** The percentage of immature (FcR+Mac-, left panel) and mature myeloid population (FcR+Mac+, right panel) of HSC liquid culture on day 6. **D)** The transcription start site (TSS) accessibility of *Ubc*Cre positive WT, Stag2, Npm1c and Stag2Npm1c mutant HSC. **E)** Volcano plot showing the chromatin accessibility of Stag2Npm1c mutant MPP3 cells comparing to WT MPP3 cells. **F)** The TSS accessibility of HSC and MPP3 comparing within each genotype.

**Supplemental Figure 8:**
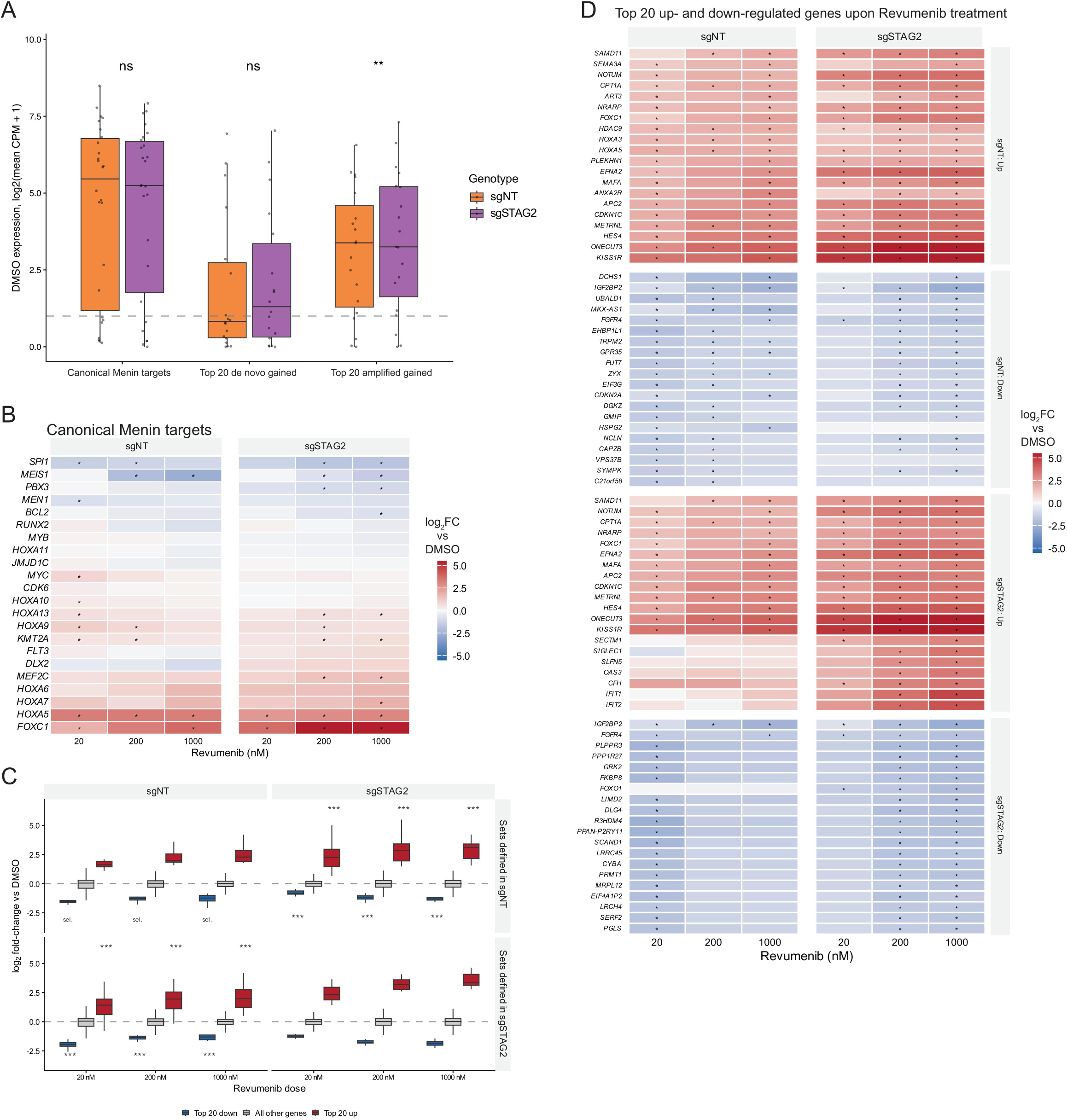
Transcriptomic response differs in sgSTAG2 OCI-AML3 cells comparing to sgNT upon Revumenib treatment. Related to figure 7. **A)** The sgNT and sgSTAG2 control gene expression level of canonical Menin targets, the top 20 de novo Menin-gained and the top 20 amplified Menin-gained in sgSTAG2. **p 0.01, paired Wilcoxon testing with Benjamini–Hochberg FDR correction. Dashed line represents CPM of 1. **B)** Heatmap of canonical Menin target expression changes in sgNT and sgSTAG2, upon Revumenib treatment at 20nM, 200nM and 1000nM doses for 48 hrs. The gene list is ranked by the Log2 fold-change. *padj 0.05, DESeq2 Wald test with Benjamini–Hochberg FDR correction. **C)** Boxplot showing the Log2 fold-change of the top 20 upregulated and top 20 downregulated gene sets defined by sgNT or sgSTAG2, upon Revumenib treatment at 20nM, 200nM and 1000nM doses for 48 hrs. ***p < 0.001, two-sided Wilcoxon rank-sum test. **D)** Heatmap of gene expression changes at the top 20 upregulated and top 20 downregulated gene set in both sgNT and sgSTAG2, upon Revumenib treatment at 20nM, 200nM and 1000nM doses for 48 hrs. The gene list is ranked by the Log2 fold-change. *padj 0.05, DESeq2 Wald test with Benjamini–Hochberg FDR correction.

